# Estimating individuals’ genetic and non-genetic effects underlying infectious disease transmission from temporal epidemic data

**DOI:** 10.1101/618363

**Authors:** Christopher M. Pooley, Glenn Marion, Stephen C. Bishop, Richard I. Bailey, Andrea B. Doeschl-Wilson

**Affiliations:** The Roslin Institute, The University of Edinburgh, Midlothian, EH25 9RG, UK; Biomathematics and Statistics Scotland, James Clerk Maxwell Building, The King’s Buildings, Peter Guthrie Tait Road, Edinburgh, EH9 3FD, UK

## Abstract

Individuals differ widely in their contribution to the spread of infection within and across populations. Three key epidemiological host traits affect infectious disease spread: susceptibility (propensity to acquire infection), infectivity (propensity to transmit infection to others) and recoverability (propensity to recover quickly). Interventions aiming to reduce disease spread may target improvement in any one of these traits, but the necessary statistical methods for obtaining risk estimates are lacking. In this paper we introduce a novel software tool called *SIRE* (standing for “Susceptibility, Infectivity and Recoverability Estimation”), which allows simultaneous estimation of the genetic effect of a single nucleotide polymorphism (SNP), as well as non-genetic influences on these three unobservable host traits. SIRE implements a flexible Bayesian algorithm which accommodates a wide range of disease surveillance data comprising any combination of recorded individual infection and/or recovery times, or disease status measurements. Different genetic and non-genetic regulations and data scenarios (representing realistic recording schemes) were simulated to validate SIRE and to assess their impact on the precision, accuracy and bias of parameter estimates. This analysis revealed that with few exceptions, SIRE provides unbiased, accurate parameter estimates associated with all three host traits. For most scenarios, SNP effects associated with recoverability can be estimated with highest precision, followed by susceptibility. For infectivity, many epidemics with few individuals give substantially more statistical power to identify SNP effects than the reverse. Importantly, precise estimates of SNP and other effects could be obtained even in the case of incomplete, censored and relatively infrequent measurements of individuals’ infection or survival status, albeit requiring more individuals to yield equivalent precision. SIRE represents a new tool for analysing a wide range of experimental and field disease data with the aim of discovering and validating SNPs and other factors controlling infectious disease transmission.

## 1 Introduction

In the era of rapid expansion in the human population resulting in increasing demands on food security, effective solutions that reduce the spread of infectious diseases not only in humans, but also in plants and livestock, are urgently needed. Failure of stringent biosecurity measures [2, 3] and emergence of anti-microbial resistance [4, 5] and escape mutants to viral vaccines [6, 7] demonstrate that infectious diseases cannot be combatted by conventional biosecurity and pharmaceutical interventions alone.

The advent of genome wide high density single-nucleotide polymorphism (SNP) chip panels has already led to a remarkable range of discoveries regarding the genetic regulation and biology of diseases and translation towards innovative therapeutics [8]. In agriculture, SNP chip panels have revolutionized breeding practices by facilitating genomic selection [9, 10]. In the infectious disease context genomic selection may effectively prevent or reduce disease spread by providing a means to identify and select against individuals with high genetic risk of becoming infected or transmitting infections purely based on their genetic make-up, without the need of exposing them to infectious pathogens [11]. However, to date the full host genetic basis underlying infectious disease transmission is still poorly understood.

Epidemiological models are widely used to identify risk factors for disease spread in populations. Indeed, modelling disease transmission in genetically heterogeneous populations is well established (see *e.g*.[12, 13]). Particularly relevant are so-called compartmental models in which individuals are classified as, for example, susceptible to infection, infected and infectious, or recovered (or alternatively dead). Transitions between these states are determined by three key individual traits: *susceptibility*, the relative risk of an uninfected individual to become infected when exposed to a typical infectious individual or infectious material excreted from such an individual, *infectivity*, the propensity of an individual, once infected, to transmit infection to a typical (average) susceptible individual, and *recoverability*, the propensity of an individual, once infected, to recover or die) [14, 15]. As demonstrated by numerous simulation studies, host genetic variation in any one of these traits, if correctly identified, could be exploited to reduce infectious disease spread within and across populations [15–18]. However, despite their strong epidemiological importance, the genetic regulation and co-regulation of these three host traits is largely unexplored. Whereas a plethora of studies have identified substantial heritable variation and SNPs associated with host susceptibility [18], remarkably little is known about the genetic regulation of host recoverability and infectivity, despite emerging evidence that genetic variation in these traits exists [19, 20]. In particular, it is currently not known to what extent infectivity is genetically controlled, despite compelling evidence that super-spreaders, defined as a small proportion of individuals responsible for a disproportionally large number of transmissions, are a common phenomenon in epidemics [21–23]. This shortcoming is largely because appropriate statistical methods for estimating genetic and also non-genetic (treatment) effects for all three key epidemiological traits controlling disease transmission from infectious disease data are currently lacking.

In many conventional genome-wide association studies (GWAS) [24], target traits for genetic improvement are measured directly, so establishing genetic associations is relatively straightforward. In the epidemiological setting, however, the susceptibility, infectivity and recoverability of individuals are not measured directly. Rather their effects are manifested in the infection and recovery times of individuals in the epidemic (or epidemics) as a whole. Furthermore, most conventional GWAS assume that an individual’s infection status is controlled by its own genetic susceptibility and environmental effects. From an epidemiological viewpoint however, an individual’s disease phenotype (*e.g.* infected or not) may not only depend on its own susceptibility and recoverability genes, but also on the infectiousness of other individuals in the same contact group, *i.e.* their infectivity and recoverability genes [25]. This complex interdependence between underlying and observable traits poses challenges for existing methods.

The motivation behind this paper is to introduce new statistical and computational methods that utilise information derived from observation of epidemics and trait interdependence to estimate, for the first time, genetic and other systematic effects for all three underlying epidemiological host traits. This requires combining statistical, epidemiological and genetic modelling principles. Analysis of incomplete epidemic data to draw inferences on epidemiological parameters is well established [26, 27]. However, analysing such data to draw joint inferences on both the disease epidemiology and host genetic variation has proven challenging [25, 28]. Recent studies have expanded conventional quantitative genetics threshold models to enable joint genetic evaluation of cattle susceptibility to, and recoverability from, mastitis [29, 30], which led to identification of novel SNPs and candidate genes associated with these traits [19]. However, because infectivity acts on group members rather than the focal individual itself, applying these technique to estimate genetic effects for infectivity is problematic.

Alternative approaches have focused on disentangling susceptibility from infectivity effects. For example, Anacleto *et al.* [31] developed a Bayesian inference approach to produce genetic risk estimates for host susceptibility and infectivity from epidemic time to infection data, assuming that susceptibility and infectivity are under polygenic control (*i.e.* they are determined by a large number of genes, each with small effect). This approach, however, does not incorporate genetic variation in recoverability, and does not estimate SNP effects. An alternative approach, based on the assumption that susceptibility and infectivity are controlled by two single bi-allelic genetic loci [32, 33], used a generalized linear model (GLM) to estimate relative allelic effects on host susceptibility and infectivity. Whilst an important contribution, this approach focused on the disease status of individuals at the end of each epidemic (*i.e.* discarding potentially useful information from the infection and recovery times themselves). It also failed to incorporate variation in recoverability, and relied on a number of simplifying assumptions which were found to produce biased estimates under certain circumstances. A variant of this approach [34], which adopted a GLM to analyse time-series data on individual disease status, illustrated the benefits of longitudinal records of individuals’ infection status for improving prediction accuracies of SNP effects, although it still relied on a number of simplifications that may compromise prediction accuracies and lead to unwanted bias. A further shortcoming of previous approaches [32–34] is that they ignore potential pleiotropic effects, *i.e.* SNPs affecting more than one epidemic trait. This seems unrealistic, since, for example, SNPs that control within host pathogen replication may also lower the risk that infection can establish, *i.e.* reduce susceptibility, and simultaneously reduce pathogen shedding and hence infectivity, and speed up recovery.

In this study we present a novel software tool called SIRE (standing for “susceptibility, infectivity and recoverability estimation”) that implements a Bayesian inference approach to simultaneously estimate the effects of a single SNP (importantly capturing any pleiotropy), together with that of other fixed effects (such as *e.g.* sex, breed or vaccination status) on host susceptibility, infectivity and recoverability from temporal epidemic data. This approach can be applied to a wide range of epidemic data, collected at the level of individuals, and accounts for different types of uncertainty in a statistically consistent way (*e.g.* censoring of data or imperfect diagnostic tests), and permits the incorporation of prior knowledge. We validate SIRE for a variety of simulated epidemic scenarios, comprising not only the ideal case in which infection and recovery / death times of each individual are known exactly, but also under more realistic scenarios in which epidemics are only partially observed.

## 2 Materials and methods

### 2.1 Data structure and the underlying genetic-epidemiological model

SIRE applies to individual-level disease data originating from one or more contact groups in which infectious disease is transmitted from infectious to susceptible individuals through contact. This data can come from well controlled disease transmission experiments or from much less well controlled field data (which may be less complete, but readily available in larger quantity).

In the context of disease transmission experiments in plants or livestock, epidemics are initiated by means of artificially infecting a proportion of “seeder” individuals which go on to transmit their infection to susceptible individuals sharing the same contact group. In field data contact groups may consist of animal herds, or any group of individuals sharing the same environment such as a pasture, pen, cage or pond, and infection is assumed to invade the group by some external, usually unknown, means (*e.g.* by the unintentional spread of infected material, or the introduction of an infected individual from elsewhere). For simplicity it is assumed that throughout the observation period groups are closed, *i.e.* no births, migrations, or transmission of disease between groups. This assumption generally holds for experimental studies and also for most common field situations, where a movement ban is imposed after disease notification [35].

The dynamic spread of disease within a contact group is modelled using a so-called SIR model [36]. Individuals are classified as being either susceptible to infection (S), infected and infectious (I), or recovered/removed/dead (R). Under the simple SIR model for homogeneous populations, the time-dependent force of infection for a susceptible individual *j* (*i.e.* the probability per unit time of becoming infected) is given by *λ*_*j*_(t) = *βI*(t), which is the product of an average transmission rate *β* and *I*(t), the number of infected individuals at time *t*. To incorporate individual-based variation in host susceptibility and infectivity, this simple expression for *λ*_*j*_(t) is replaced by an individual force of infection (see [31] for a formal derivation)

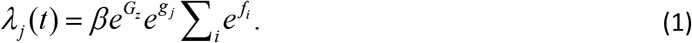

Here *g*_*j*_ characterises the fractional deviation in individual *j’s* susceptibility as compared to that of the population as a whole (*e.g. g*_*j*_=0.1 corresponds to individual *j* being ≃10% more susceptible than the population average), *f*_*i*_ characterises the corresponding quantity for individual *i’s* infectivity, and the sum in Eq.(1) goes over all individuals infected at time *t* sharing the same contact group *z* as individual *j* (note, this sum varies as a function of *t* as individuals become infected and recover). The term *G*_*z*_ in Eq.(1) accounts for the fractional deviation in disease transmission for group *z*. This incorporates group-specific factors that influence the overall speed of an epidemic in one contact group relative to another (*e.g.* animals kept in different management conditions, environmental differences, or variation in pathogen strains with differing virulence). Whilst variation in *G*_*z*_ may be small for a well-controlled challenge experiment, this may not be the case in real field data. *G*_*z*_ is assumed to be a random effect with standard deviation σ_G_. The exponential dependencies in Eq.(1) ensure that *λ*_*j*_ is strictly positive and allow for the possibility that some groups or individuals are much more/less susceptible/infectious than others, *i.e.* it can accommodate potential super-spreaders.

Whilst in Eq. (1) infection is modelled as a Poisson process with individual infection rates *λ*_*j*_ [18, 20], the recovery process is modelled by assuming that the time taken for individual *m* to recover after being infected is drawn from a gamma distribution with an individual-based mean *w*_*m*_ and shape parameter *k* (which for simplicity is assumed to be the same across individuals). This mean recovery time is expressed as

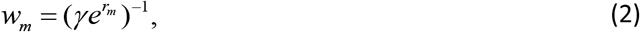

where γ represents an average recovery rate across the population and *r*_*m*_ describes the fractional deviation from this for individual *m*. This approach is taken to allow the recovery probability distribution to adopt a more biologically realistic profile compared with the exponential distribution often assumed (see electronic supplementary material Appendix A for further details).

Following standard quantitative genetics theory [37], the individual-based deviations in susceptibility ***g***, infectivity ***f*** and recoverability ***r*** (which are vectors with elements relating to each individual) are decomposed into the following contributions

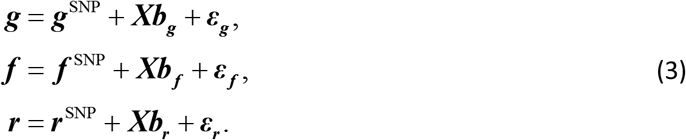

#### SNP effects

The model assumes that a specific locus defined by a SNP (potentially) plays an important contribution to the trait values (note, repeated analysis can be performed on different SNPs of interest). Assuming a diploid genomic architecture with biallelic SNP implies three SNP genotypes: *AA, AB* and *BB*. The SNP contribution to the traits for individual *j* depends on *j*’s genotype in the following way:

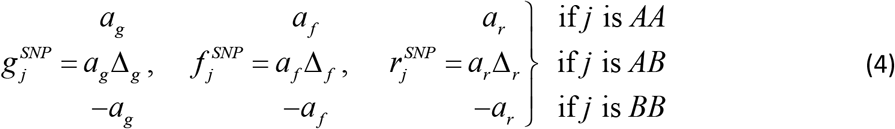

The parameters *a*_*g*_, *a*_*f*_ and *a*_*r*_ capture the relative differences in trait values between *AA* and *BB* individuals, and are subsequently referred to as the “SNP effects” for susceptibility, infectivity and recoverability, respectively (*e.g.* if *a*_*g*_ is positive, individuals with an *AA* genotype will, on average, be more susceptible to disease than those with a *BB* genotype). The scaled dominance factors Δ_*g*_, Δ_*f*_ and Δ_*r*_ characterise the trait deviations between the heterozygote *AB* individuals and the homozygote mean (a value of 1 corresponds to complete dominance of the *A* allele over the *B* allele and −1 when the reverse is true, whereas absence of dominance is represented by a value of 0) [38].

#### Fixed effects

The design matrix ***X*** and fixed effect vectors ***b***_***g***_, ***b***_***f***_ and ***b***_***r***_ in Eq.(3) allow for other known sources of variation to be accounted for (*e.g.* breed, sex or vaccination status). Following convention, an additional fixed effect is added to account for trait mean, which is explicitly chosen to ensure the population averages of ***g***, ***f*** and ***r*** are zero (remembering that the average effects are already captured by the parameters β and γ).

#### Residual contributions

Here **ε**=(**ε**_**g**_, **ε**_**f**_, **ε**_**r**_) accounts for all other contributions to the traits (*i.e.* coming from genetic effects not captured by the SNP under consideration, as well as any non-genetic environmental variation). We assume that for each individual the three trait residuals are drawn from a single multivariate normal distribution with zero mean and 3×3 covariance matrix **Σ**.

Including these correlations is important because it allows for the possibility that, for example, more susceptible individuals may also, on average, be more infectious and recover at a slower rate (on top of any correlations which may also arise from the SNP and fixed effects). Note that in this study, which focuses on the estimation of SNP effects, there is no explicit distinction between random genetic and environmental effects, although the model could be extended to incorporate estimation of these polygenic effects. It is thus assumed that individuals are randomly distributed across the groups with respect to the genetic effects on the epidemiological traits not captured by the SNP. Also note that Eq.(3) does not contain random group effects for the individual epidemiological traits. This is because the group effect has already been incorporated in the expression of the individual force of infection in Eq.(1). In other words, it is assumed that the group environment is the dominant mechanism affecting the speed at which infection spreads within a group rather than group specific factors affecting individuals’ susceptibility, infectivity or recoverability.

### 2.2 Bayesian inference

Based on the description above, the model contains the following set of parameters: θ=(*β*, *γ*, *k*, *a*_*g*_, *a*_*f*_, *a*_*r*_, Δ_*g*_, Δ_*f*_, Δ_*r*_, ***b***_***g***_, ***b***_***f***_, ***b***_***r***_, **ε**_***g***_, **ε**_***f***_, **ε**_***r***_,, **Σ**, ***G***, σ_G_). We denote the complete set of infection and recovery event times for all individuals as ξ over the observed duration of the epidemics [39]. Typically ξ is not precisely known, and so we consider the general case in which ξ represents a set of latent model variables. The nature of the actual observed data *y* will be problem dependant. For example, in some instances recovery or removal (*e.g.* due to death) times will be precisely known but infection times completely unknown. In other instances infection and recovery times will both be unknown, but results from disease diagnostic tests provide information regarding disease status at particular points in time. The framework presented in this paper is flexible to these various possibilities.

Application of Bayes’ theorem implies that the posterior probability distribution for model parameters and latent variables is given by

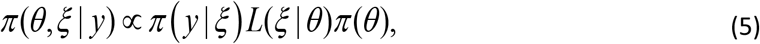

where individual components are defined as follows:

**Observation model *π*(*y*|ξ)** – the probability of the data given a set of event times ξ. The expression for the observation model depends on the nature of the data being observed. In many contexts this simply takes the values one or zero depending on whether ξ is consistent with *y* or not. For example a perfect disease diagnostic test showing that an individual is infected would be only consistent with ξ containing an infection event on that individual *prior* to the time of the test and a recovery event *after* the time of the test. Similarly, if data *y* indicates that an individual becomes infected at a particular point in time, this is only consistent provided ξ also contains this infection event. When imperfect disease diagnostic test results are available the observation model includes the sensitivity and specificity of the test to account for this uncertainty in the data. In summary, the observation model depends on the data collection process and constrains the possible event sequences ξ, and this, in turn, informs the model parameters θ.
**Latent process likelihood *L*(ξ|θ)** – the probability of ξ being sampled from the model given parameters θ. This can be derived from the genetic-epidemiological model described in the previous section [26, 27] (see Appendix B for details), and is given by

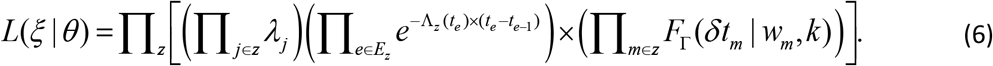 The functional dependence of *L*(ξ|θ) on the parameters θ is expressed in terms of the force of infections λ_*j*_ in Eq.(1) and mean recovery times *w*_*m*_ in Eq.(2), which themselves depend in ***g***, ***f*** and ***r*** in Eq.(3). The product *z* goes over all contact groups and within each contact group: *j* goes over individuals that become infected *excluding* those which initiate epidemics [40], *m* goes over individuals that become infected *including* those which initiate epidemics and *e* goes over both infection and recovery events (with corresponding event times *t*_*e*_). Here the notation *j*∈*z* indicates that *j* goes over all those individuals *j* in contact group *z*, and *e*∈*E*_*z*_ indicates that *e* goes over all events *E*_*z*_. The force of infection λ_*j*_ is calculated immediately prior to individual *j* becoming infected. The gamma distributed probability density function *F*_*Γ*_ for recovery events gives the probability an individual is infected for duration *δt*_*m*_ given a mean duration *w*_*m*_ and shape parameter *k*. The time dependent total rate of infection events Λ_z_ in contact group *z* immediately prior to event time *t*_*e*_ is given by

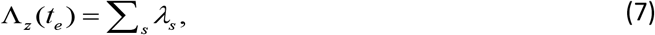

where the sum in *s* goes over all susceptible individuals in group *z* at that time. An important point to mention is that Eq.(6) is calculated on an unbounded time line. In situations in which data is censored, the observation model restricts events that occur within the observed time window, but other events can exist outside of this observed region [41].
**Prior*π*(θ)** – the state of knowledge prior to data *y* being considered. To account for the prior assumption that residuals ***ε*** in Eq.(3) are multivariate normally distributed and that the vector of group effects ***G*** in Eq.(1) are random effects, *π*(θ) can be decomposed into

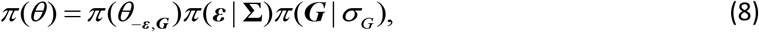

where θ_−***ε,G***_ includes all parameters with the exception of ***ε*** and ***G*** and

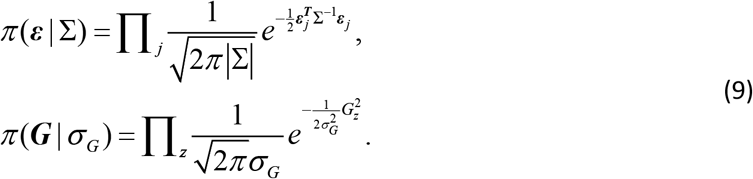

Here *j* goes over each individual and **ε**_***j***_ =(ε_*g,j*_, ε_*f,j*_, ε_*r,j*_)^T^ is a three dimensional vector giving the residual contributions to the susceptibility, infectivity and recoverability of *j*. **Σ** is a 3×3 covariance matrix (which describes not only the overall magnitude of the residual contributions, but also any potential correlations between traits). Finally, the product *z* in Eq.(9) goes over all contact groups and *G*_*z*_ represents the group-based fractional deviation in transmission rate, which is assumed to be independent between groups and normally distributed with standard deviation σ_G_.

The default prior for θ_−***ε,G***_ (which can be modified if necessary) is largely uninformative but does place upper and lower bounds on many of the key parameters to stop them straying into biologically unrealistic values (details are given in Appendix C).

Samples for θ and ξ from the posterior are generated by means of an adaptive Markov Chain Monte Carlo (MCMC) schemes which implements optimised random walk Metropolis-Hastings updates for most parameters and posterior-based proposals [1] to aid fast mixing of the residual parameters (details are given in Appendix D).

### 2.3 SIRE

SIRE is a desktop application that implements the Bayesian algorithm outlined above. It is freely available to download from the supplementary material or at www.mkodb.roslin.ed.ac.uk/EAT/SIRE.html (with versions for Windows, Linux and Mac). An easy to use point and click interface allows for data tables to be imported in a variety of formats and graphical outputs are dynamically displayed as inference is performed. The core of SIRE utilises efficient C++ code and allows for running MCMC chains on multiple CPU cores.

SIRE takes as input any combination of information about infection times, recovery times, disease status measurements, disease diagnostic test results, genotypes of SNPs or any other fixed effects (see screenshot in Fig 1a), details of which individuals belong to which contact groups and any prior specifications (Fig 1b). The output from SIRE consists of posterior trace plots for model parameters θ, distributions (Fig 1c), visualisation of infection and recovery times ξ, dynamic population estimates and summary statistics (means and 95% credible intervals) as well as MCMC diagnostic statistics (Fig 1d). Posterior distribution graphs can be exported from SIRE and also files containing posterior samples of θ and ξ for further analysis using other tools. The user guide for SIRE is available in the electronic supplementary material and on the website.

**Fig 1.**
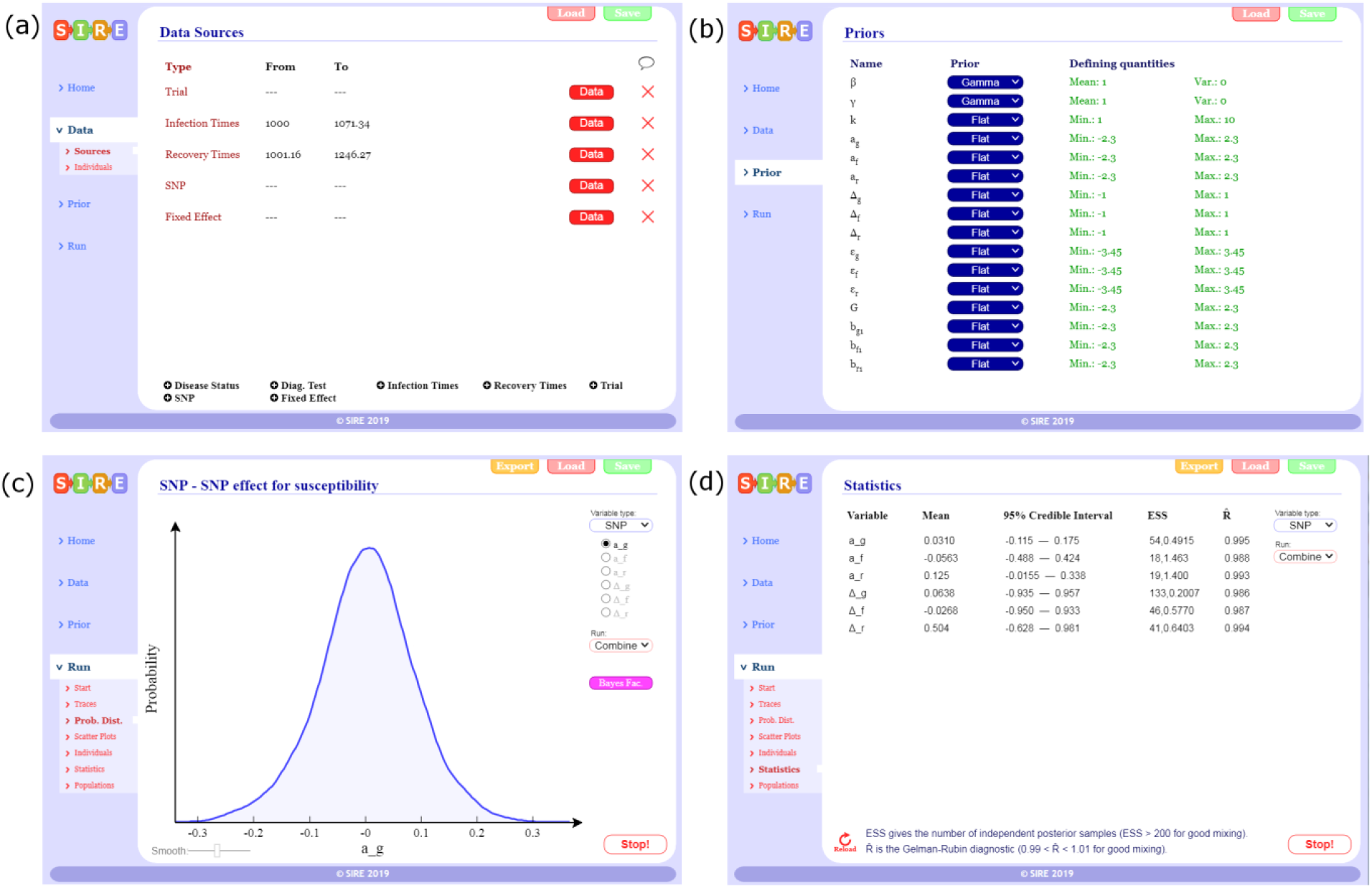
SIRE software. Illustrative screenshots of the software package: (a) Different data sources can be imported by loading user defined data tables (text or cvs files), (b) prior specification can be made on parameters, (c) posterior distributions can be visualised as inference in being performed, and (d) summary statistics and MCMC diagnostics.

### 2.4 Data scenarios

SIRE is flexible to many possible inputs. Reflecting real-world datasets this paper considers five potential data scenarios (DS):

#### DS1: Infection and recovery times for all individuals exactly known

This represents the best case scenario for inferring parameter values. For example, appearance of symptoms or visual or behavioural signs may indicate the onset of infection, and recovery/removal times are given by the time of death.

#### DS2: Only recovery times known

Often “recovery” in compartmental SIR models represents the death and removal of individuals. Consequently DS2 is pertinent to cases in which the only measurable quantity is the time at which individuals die. For example, disease challenge experiments in aquaculture routinely record time of death rather than infection times, which are usually difficult to measure [42].

#### DS3: Only infection times known

Whilst less common than DS2, in some instances data provides information regarding when individuals become infected but not when they recover. For example in human epidemics, patients may go to the doctor when they become ill, but no records will be kept on when they recover.

#### DS4: Disease status periodically checked

DS4 represents the most common scenario for monitoring infectious disease spread in livestock or plant populations, where each individual is periodically checked to establish its disease status. Under DS4 the point at which epidemics start is usually unknown, as well as the infection and recovery times of individuals themselves. However the diagnostic test results place constraints on these quantities. For example, if an individual is found to be uninfected at one sampling time and infected at the next sampling time this means that infection must have occurred at some point in the intervening period (note here we assume perfect diagnostic tests but SIRE also allows for imperfect diagnostic test results to be used, provided the sensitivity and specificity of the tests are known).

#### DS5: Time censored data

This data scenario relates to situations in which epidemics are not observed over their entire time period. For example a disease transmission experiment being carried out may be terminated early, due to cost or other factors (*e.g.* animal welfare), even though epidemics have not completely died out.

## 3 Assessment of performance and data requirements

In this section we apply SIRE to simulated datasets in order to 1) test the extent to which the inferred posterior parameter distributions agree with their true values, and 2) investigate how the precision, accuracy and bias of inferred model parameters depends on the type of data available.

Initially the focus of results will be on DS1 (which although rarely applies in practice, still provides useful insights for software validation and application) and later in section 3.5 consideration is given to DS2-5.

### 3.1 Illustrative example simulation and inference

We first demonstrate the performance of SIRE assuming complete information of individuals’ infection and recovery times, for a representative but complex set of parameters with regards to the genetic and non-genetic regulation of the three epidemiological host traits. Subsequently we investigate how these results change under different parameter and data scenarios.

#### Simulations

Individuals were randomly assigned into *N*_*group*_ different contact groups, with each group containing *G*_*size*_ individuals. The SNP under investigation was assumed to be in Hardy-Weinberg equilibrium [38] with an *A* allele frequency of *p*=0.3. For the effect sizes we used the values *a*_*g*_=0.4, *a*_*f*_=0.3, *a*_*r*_=−0.4, representing a relatively large pleiotropic effect (which confers higher susceptibility for *AA* compared to *BB* individuals, as well as slightly higher infectivity and reduced recoverability). The choice of Δ_*g*_=0.4, Δ_*f*_=0.1, Δ_*r*_=−0.3 for the scaled dominance factors represents partial, but not strong, dominance of either the *A* or *B* allele. For simplicity we included only a single fixed effect, *e.g.* sex, of arbitrary moderate size *b*_*g0*_=0.2, *b*_*f0*_=0.3, *b*_*r0*_=−0.2 with individuals in the population randomly selected to be male or female. The residual variances were chosen to be Σ_*gg*_=Σ_*ff*_=Σ_*rr*_=1, corresponding to a large variation in traits between individuals (perhaps larger than is biologically realistic, but here we want to demonstrate that inference of the SNP effects is still possible *despite* significant variation in trait values arising from other sources). In line with the direction of the SNP effects, the covariances were chosen to be Σ_*gf*_=0.3, Σ_*gr*_=−0.4 and Σ_*fr*_=−0.2, representing a potential scenario in which individuals that are more susceptible are also more infectious and recover at a slower rate and *vice-versa*). To accommodate variation in epidemic speed across groups, we set the standard deviation in the group effects to σ_G_=0.5. Finally, the average transmission rate was chosen to be β=0.3/*G*_*size*_ (selected because it led to a substantial fraction of individuals becoming infected and including *G*_*size*_ such that the basic reproductive ratio *R*_*0*_ remained independent of group size, *i.e.* frequency dependent transmission) and an average recovery rate γ=0.1 with shape parameter *k=*5 (corresponding to the infection duration being relatively highly peaked around a mean of 10 time units).

Simulated epidemic data was generated by means of a Doob-Gillespie algorithm [43] modified to account for non-Markovian recovery times (details of this procedure are given in Appendix F). A typical output for one simulated epidemic in a single contact group *N*_*group*_=1 with *G*_*size*_=50 individuals is shown in Fig 2. Whilst the simulation itself is generated on an individual basis, this graph summarises dynamic variation in the susceptible, infectious and recovered populations, categorised by SNP genotype. It reveals classic epidemic SIR model behaviour: a single infected individual passes its infection on to others, triggering a rapidly spreading infection process throughout the population until the epidemic eventually dies out as a result of the susceptible population becoming largely exhausted and the remaining infected population recovering. Note that in closed groups not all susceptible individuals become infected. In this particular case some *AB* and *BB* individuals remain uninfected at the end of the epidemic. The absence of *AA* individuals partly stems from natural stochasticity in the system, but also partly from the fact that *a*_*g*_=0.4 is positive, *i.e. AA* individuals are more susceptible to disease and so on average less likely to remain uninfected. Consequently we can link the genetic composition in the final state of the epidemic to the expected value for *a*_*g*_ (which, based on this particular dataset, is more likely positive than negative). Over and above information from the final state, however, there is much to be gained from also accounting for the infection and recovery event times themselves. The Bayesian approach adopted in this paper utilises all this information to extract the best available parameter estimates.

**Fig 2.**
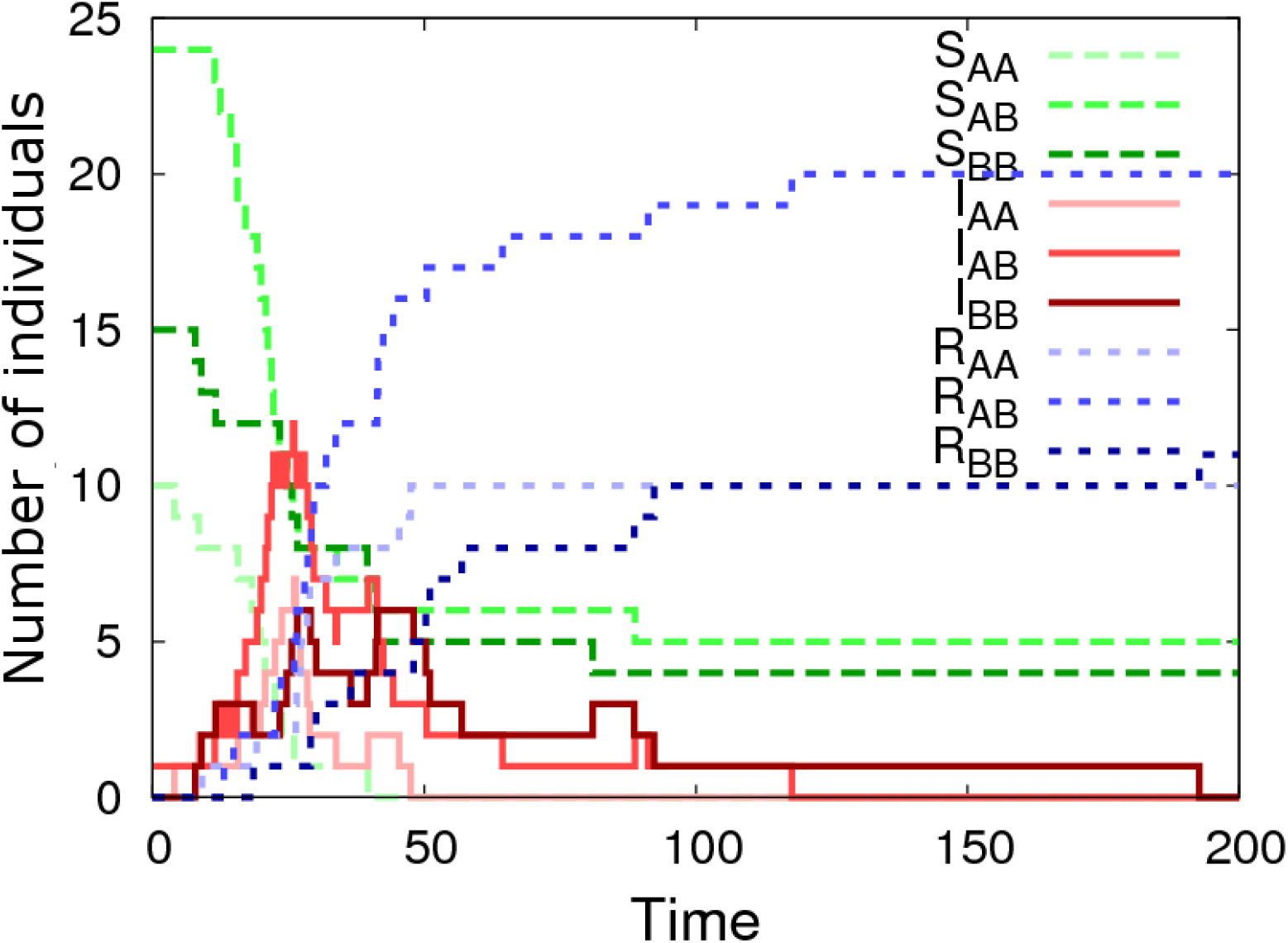
Simulated epidemic profiles. This graph shows epidemic profiles for the three SNP genotypes (*i.e. AA*, *AB* or *BB*), where S_*g*_, I_*g*_, R_*g*_ indicate the number of susceptible, infected and recovered individuals of genotype *g*, respectively. This example is simulated using a single contact group containing *G*_*size*_=50 individuals, of which one is initially infected. The model parameters θ are: β=0.006, γ=0.1, *k=*5, *a*_*g*_=0.4, *a*_*f*_=0.3, *a*_*r*_=−0.4, Δ_*g*_=0.4, Δ_*f*_=0.1, Δ_*r*_=−0.3, *b*_*g0*_=0.2, *b*_*f0*_=0.3, *b*_*r0*_=−0.2, Σ_*gg*_=1, Σ_*gf*_=0.3, Σ_*gr*_=−0.4, Σ_*ff*_=1, Σ_*fr*_=−0.2, Σ_*rr*_=1, σ_G_=0.5 and the *A* allele has frequency *p=0.3*. Note, the step jumps in curves result from discrete disease status transitions in individuals.

The information content from a single epidemic is generally insufficient to estimate the large number of parameters in the model. Therefore we next simulated a more realistic dataset (using the same parameter set as above) made up of 1000 individuals split into *N*_*group*_=20 contact groups, each containing *G*_*size*_=50 individuals. The infection and recovery event times from this simulation were then used as input data into SIRE (scenarios in which infection and recovery times are not known precisely are discussed later in section 3.5).

#### Parameter estimates

Fig 3 shows the inferred posterior probability distributions for all parameters in θ corresponding to the simulated multi-group scenario described above. The actual parameter values used to generate the data (see vertical black dashed lines in Fig 3) consistently lie within regions of high posterior probability. The standard deviations (SDs) in these distributions characterise the precision with which parameters can be estimated:

**Population average parameters (Figs 3a-c)** – The recovery rate γ has the greatest precision (smallest relative SD), followed by the transmission rate β. Whilst the distribution for the shape parameter *k* is wide, it is clearly able to discount the possibility of an exponential recovery duration (*i.e.* k=1), which has a very low posterior probability, over a more peaked distribution (*i.e*. *k*>1).
**SNP effects (Figs 3d-f)** – The estimated recovery SNP effect *a*_*r*_ is highly peaked around its true value of −0.4 (Fig 3f). Importantly this distribution has an extremely low posterior probability at *a*_*r*_=0. Indeed, since *a*_*r*_=0 does not lie within the 95% credible interval it can be concluded, to a high degree of certainty, that the SNP *is* associated with recoverability. The same is true for the susceptibility SNP effect *a*_*g*_ in Fig 3d, albeit with a wider posterior probability distribution. This difference is for two reasons: firstly the recovery process involves only *a*_*r*_, whereas the infection process involves both *a*_*g*_ and *a*_*f*_ (leading to potential confounding between these parameters) and secondly the recovery processes is gamma distributed which has a smaller standard deviation than the more dispersed Poisson process governing infection. The infectivity SNP effect *a*_*f*_ in Fig 3e exhibits a much wider probability distribution than the other two SNP effects. The fact that zero *does* lie within the 95% posterior credible interval (which goes from −0.35 to 2.1) means that no certain association with infectivity can be made in this particular example. Figs 3d-f illustrates a general principle that was common in the vast majority of subsequent simulation scenarios: SNP effects associated with recoverability are most precisely estimated, followed by susceptibility, and finally infectivity [44].
**Scaled dominance factor** (Figs 3g–i) – Compared to the SNP effects themselves, precision of the scaled dominance parameters is relatively poor, and actually reduces as the size of the SNP effects goes down (results not shown), which makes sense in the limit of zero SNP effect size, because here no information about dominance is available. Estimating them accurately, therefore, either requires very large SNP effects or substantially more data.
**Fixed effects** (Figs 3j–l) – Since SNP effects are also a type of fixed effect, the same comments as above also apply for other fixed effects.
**Residual covariance matrix and random group effect** (Figs 3m–s) – Interestingly, it was possible to obtain relatively good estimates for elements in the residual covariance matrix. Again, the familiar pattern is observed whereby quantities related to recoverability are more precisely estimated than those related to susceptibility, with infectivity the least precise. Finally, the variance of the group effect could be estimated with similar precision as that for susceptibility (Figs 3s & m).

**Fig 3.**
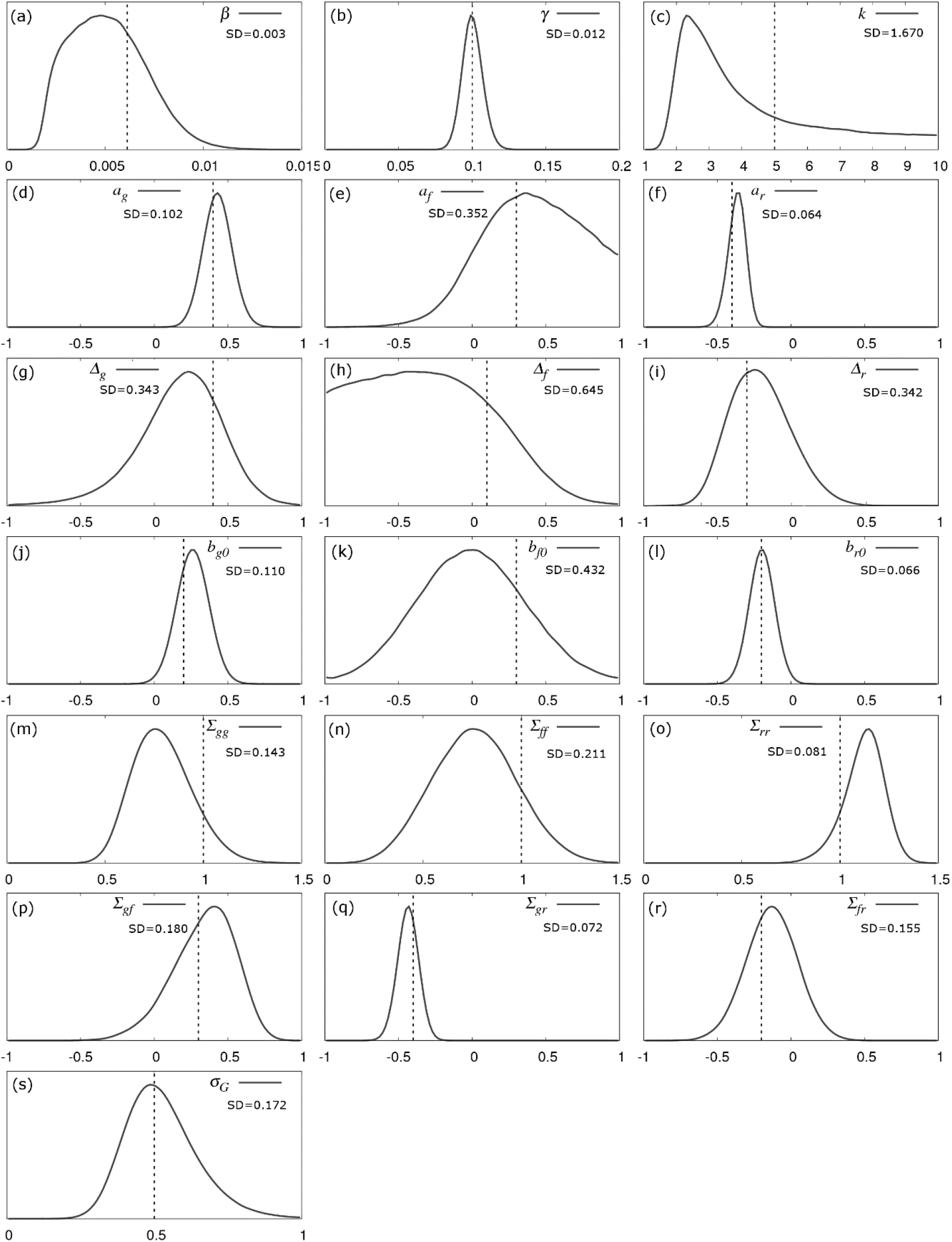
Parameter posterior distributions. Probability distributions for model parameters inferred from a simulated dataset which consisted of exact infection and recovery times (DS1) for *N*_*group*_=20 contact groups each containing *G*_*size*_=50 individuals. The parameter values in Fig 1 were used for the simulation (denoted by the vertical black dashed lines). The standard deviations (SD) give a measure of precision.

### 3.2 Dependence on parameter values

The previous section showed an illustrative example for a particular parameter set. Here we assess what happens when different parameters in the model are altered. This was achieved by means of taking the following “base” set of parameters

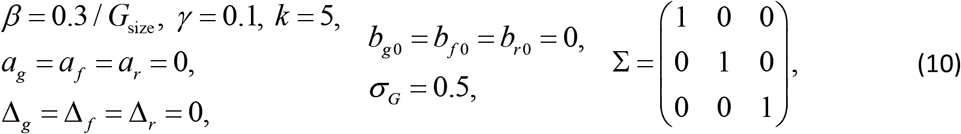

and then changing each parameter separately (fixing all others) [45]. Fig 4 shows scatter plots (each referring to a different selected parameter) of the posterior means (crosses) with corresponding 95% credible intervals inferred from a single simulated dataset using the true selected parameter value on the *x*-axis. Plots in which most crosses lie near to the diagonal line imply that inference is able to accurately capture the true parameter values. Table 1 shows the corresponding prediction accuracy, measured as the correlation between the inferred and true parameter values. Except for Δ_*f*_ for which prediction accuracy was only 34%, prediction accuracies for all other parameters ranged from 69-99%. In line with the discussion above, parameters associated with recoverability have generally higher predication accuracies than those associated with susceptibility, which are again higher than those for infectivity.

**Table 1.** Prediction accuracy, bias and precision for the parameter estimates. Other columns relate to the sub-plots in Fig 4 (see Fig 4 caption for information about the underlying data). Prediction accuracy is defined as the correlation between the inferred and true parameter values. The *y*-intercept and slope were obtained by fitting regression lines through the data points in Fig 4 (a *y*-intercept of zero and slope of one indicates no bias). Av. SD gives the average posterior standard deviation across all data points as an indicator for precision of parameter estimates. Subscripts *g*, *f* and *r* refer to susceptibility, infectivity and recovery, respectively.

| Parameter | Accuracy | y-intercept | Slope | Av. SD | Description |
| --- | --- | --- | --- | --- | --- |
| $\beta$ | 0.833 | 0.000 | 1.080 | 0.003 | Average transmission rate |
| $\gamma$ | 0.982 | 0.001 | 0.999 | 0.013 | Average recovery rate |
| $k$ | 0.806 | 1.810 | 0.633 | 1.580 | Recovery shape parameter |
| $a_g$ | 0.985 | -0.004 | 1.020 | 0.091 | SNP effect for susceptibility |
| $a_f$ | 0.875 | -0.054 | 0.860 | 0.287 | SNP effect for infectivity |
| $a_r$ | 0.995 | -0.020 | 0.990 | 0.065 | SNP effect for recoverability |
| $\Delta_g$ | 0.910 | -0.038 | 0.751 | 0.439 | Dominance factor (per trait) |
| $\Delta_f$ | 0.335 | 0.065 | 0.133 | 0.530 | |
| $\Delta_r$ | 0.920 | -0.005 | 0.781 | 0.373 | |
| $b_{g0}$ | 0.978 | -0.012 | 1.000 | 0.105 | Fixed effect (per trait) |
| $b_{f0}$ | 0.871 | -0.035 | 1.100 | 0.365 | |
| $b_{r0}$ | 0.992 | 0.008 | 1.000 | 0.073 | |
| $\Sigma_{gg}$ | 0.885 | 0.101 | 0.903 | 0.136 | Residual covariance matrix |
| $\Sigma_{ff}$ | 0.691 | 0.264 | 0.563 | 0.203 | |
| $\Sigma_{rr}$ | 0.981 | 0.027 | 1.000 | 0.071 | |
| $\Sigma_{gf}$ | 0.789 | -0.022 | 0.949 | 0.230 | |
| $\Sigma_{gr}$ | 0.978 | 0.000 | 0.959 | 0.067 | |
| $\Sigma_{fr}$ | 0.862 | 0.002 | 0.983 | 0.144 | |
| $\sigma_G$ | 0.899 | 0.008 | 1.071 | 0.144 | SD of group effects |

**Fig 4.**
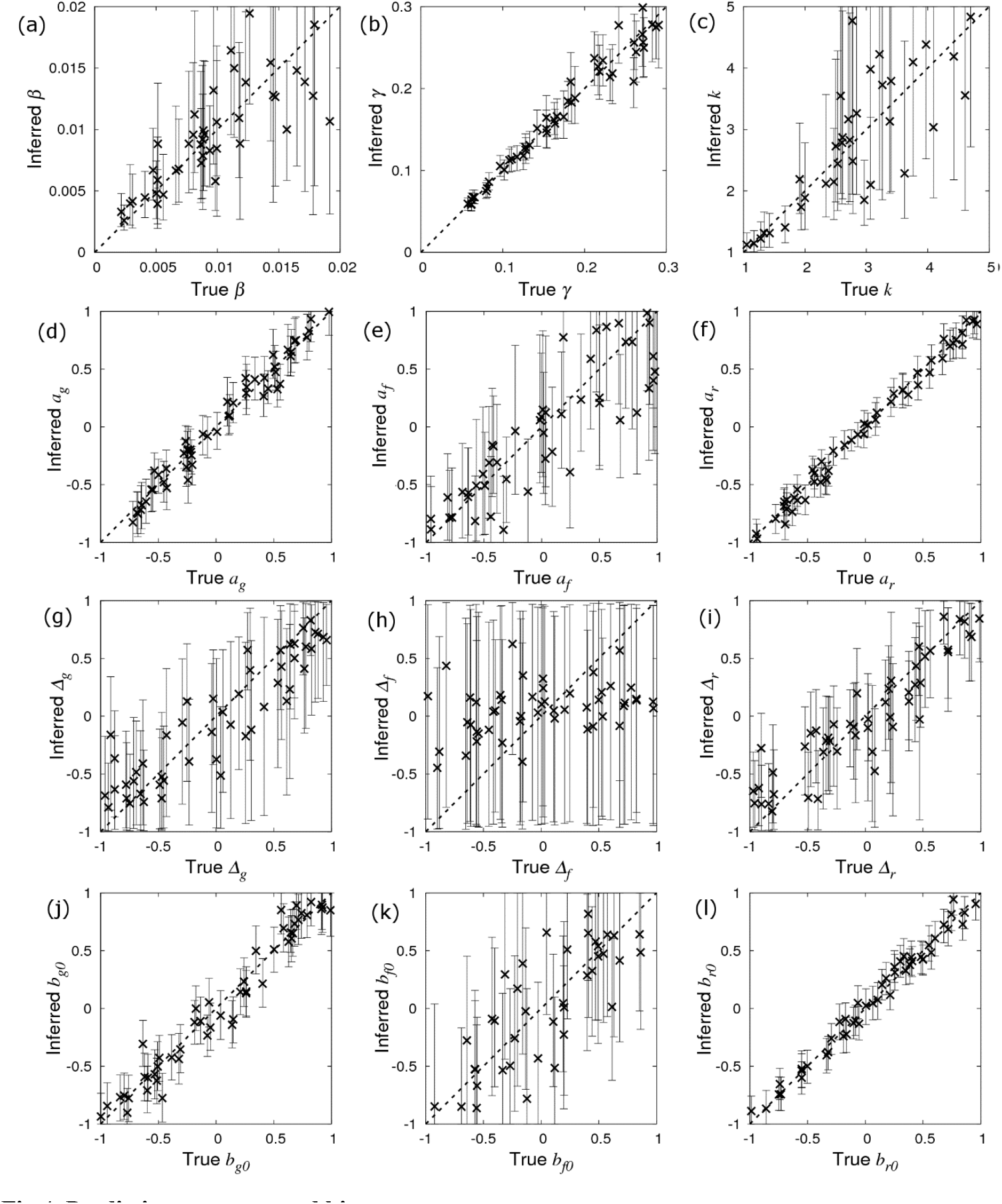
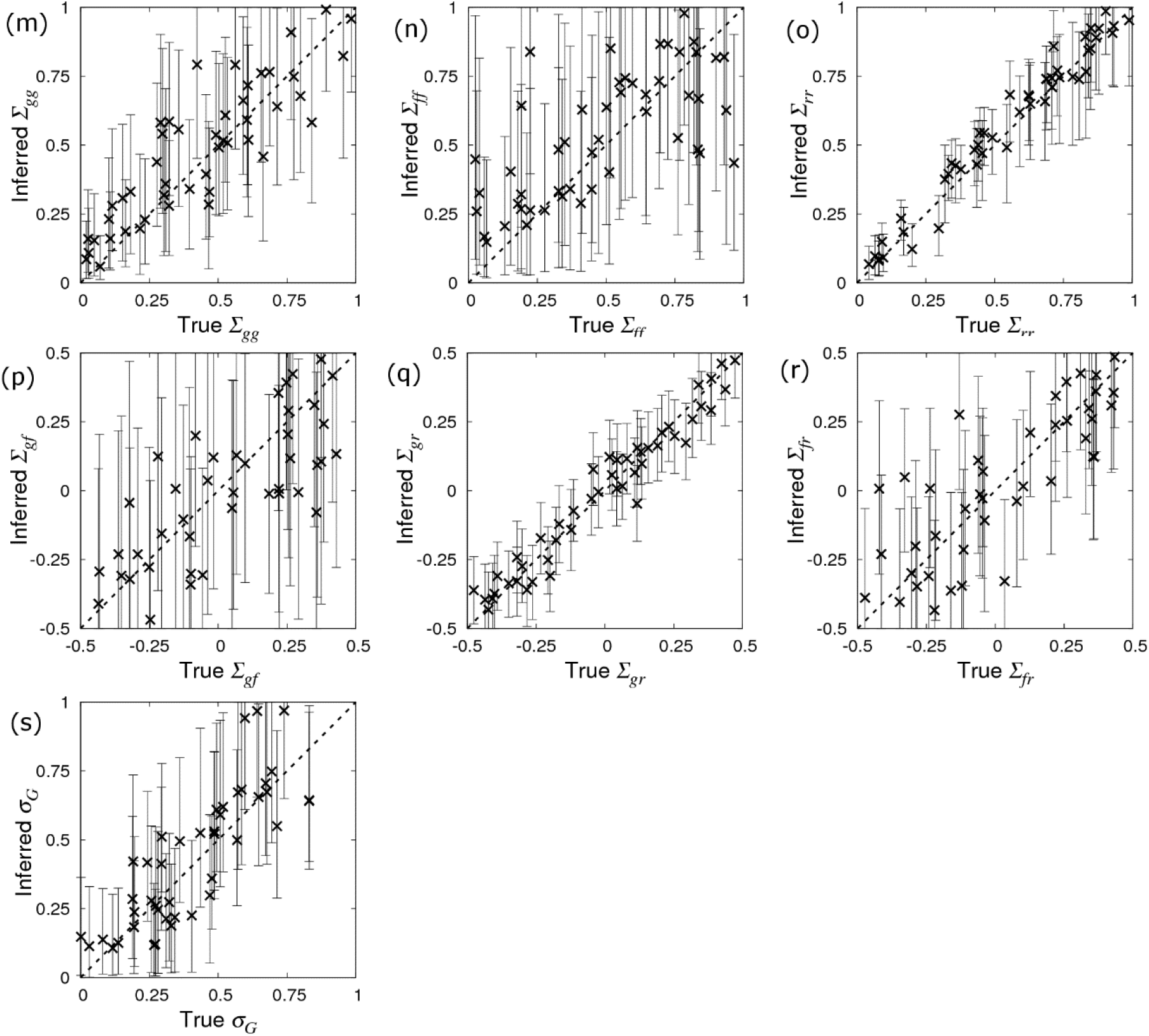
Prediction accuracy and bias. The inferred posterior distributions for parameters compared to their true value. Simulated data was generated using the base parameter set in Eq.(10) except for a single parameter which was singled out in each of the sub-plots above*. Each cross corresponds to the inferred posterior mean (with error bars indicating 95% credible intervals) of the selected parameter (whose true value is on the *x*-axis) when SIRE is applied to a single simulated dataset consisting of infection and recovery times (DS1) from *N*_*group*_=20 contact groups each containing *G*_*size*_=50 individuals. A description of the model parameters, together with calculated prediction accuracies (correlation between true and inferred value), and bias (represented by intercept and slope of regression lines fitted to the data points), and average standard deviations are given in Table 1. (*Additionally for (g) *a*_*g*_=0.4, (h) *a*_*f*_=0.4 and (i) *a*_*r*_=0.4, such that dominance has an effect).

Bias indicates systematic differences between the true parameter values and those inferred from the data. Bias was measured by fitting regression lines through the posterior means in Fig 4 (as a function of the true parameter value). The corresponding *y*-intercept and slope values are shown in Table 1, where a zero *y*-intercept and a slope of one indicate absence of bias. Whilst the majority of observed *y*-intercepts tended to be very small, the slope for some of the parameters is markedly less than one (most notably for Δ_*f*_). The reason for this is as follows. When Bayesian analysis reveals insufficient information regarding a parameter, its distribution follows that of the prior (which are uniform for all the parameters in this particular study, as described in Appendix C). This behaviour happens irrespective of the parameter’s true value, leading to a plot in Fig 4 that would be entirely flat (*i.e.* a slope of zero). Therefore, the slopes of less than one in Fig 4 simply reflect a lack of data, which is essentially another manifestation of a lack of parameter precision. Consequently, bias reduces as the amount of data increases (provided the model being fitted is the correct one).

From the point of view of this paper, the probability distributions which are of greatest interest are the SNP effects. Noting the sizes of the error bars across Figs 4d-f demonstrate that the precisions of the parameter estimates are largely independent of the values of the parameters themselves, a result which can be supported analytically [46]. This implies that the precision of SNP effects calculated using the base set of parameters in Eq.(10) is expected to be generally applicable to any other parameter set [47] (*e.g.* the average SDs in Table 1 for the base parameter set are very similar to the SDs shown in Fig 3).

Consequently, the remainder of this paper focuses on investigating how SNP effect estimates are affected by contact group structure and the nature of the measured data using this base set of parameters. We focus first on outlining the behaviour with respect to key design features, *e.g.* group size, number of individuals per group and allele frequency, and then go on to consider how observations of the system influence what can be learned.

### 3.3 Dependence on the number and size of contact groups

The crosses in Fig 5 shows how SDs in the SNP effects change as a function of the number of individuals *G*_*size*_ within each contact group (here *N*_*group*_=10 contact groups are assumed). The SD in *a*_*g*_ reduces as the number of individuals in each contact group *G*_*size*_ increases (Fig 5a). Importantly this relationship scales as a line of slope −½ (note the log scales on this plot), corresponding to precision increasing by a factor of two as the number of individuals is increases by a factor of four (in line with what would be expected from central limit theorem). Fig 5a provides insights into how many individuals would need to be observed in order to be able to make an association with a susceptibility SNP effect of a given size. For example, in order to detect an association with a susceptibility SNP of effect size *a*_*g*_ = 0.4, *G*_*size*_=20 individuals per contact group, and so *G*_*size*_×*N*_*group*_=200 individuals in total would be needed to assure that the 95% credibility interval does not contain zero (assuming approximate normality for the posterior distribution), as illustrated by that black dashed line in Fig 5a. Fig 5c shows the same scaling relationship for identifying recoverability SNP effects, but this time only *G*_*size*_×*N*_*group*_=100 individuals are needed to make associations for recovery SNP effects (reflecting the fact that *a*_*r*_ can be inferred more precisely, as mentioned previously). A very different state of affairs, however, is observed in Fig 5b. Here we see that not only is the infectivity SNP effect *a*_*f*_ poorly estimated, but also its precision does not markedly improve even when the number of individuals in each contact group *G*_*size*_ is substantially increased.

**Fig 5.**
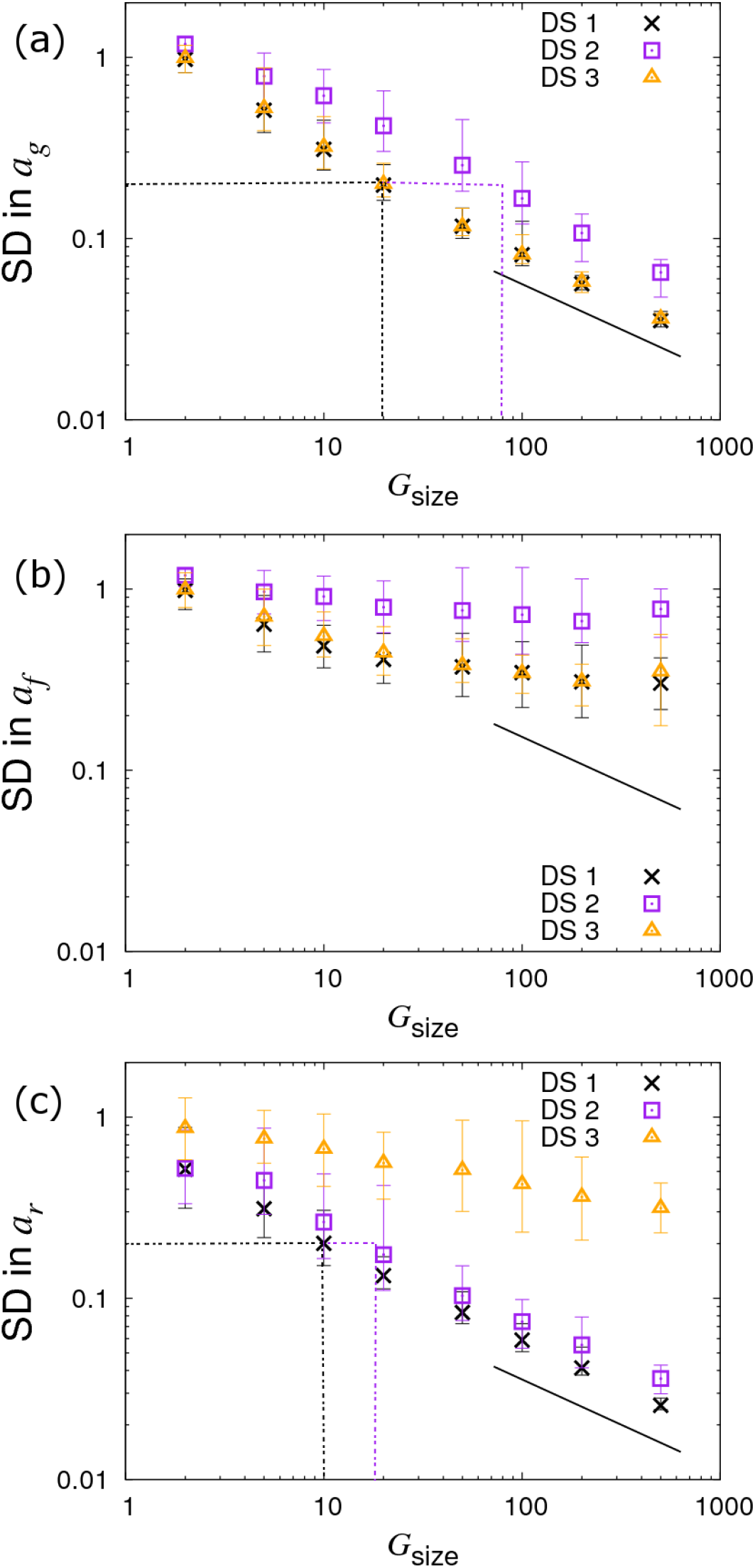
Variation in precision of the SNP effect estimates with group size *G*_size_. Posterior standard deviations (SDs) in SNP effects for (a) susceptibility *a*_*g*_, (b) infectivity *a*_*f*_ and (c) recoverability *a*_*r*_ from simulated data with *N*_*group*_=10 contact groups each containing *G*_*size*_ individuals (which is varied). Different symbols represent different data scenarios: DS1) Both the infection and recovery times for individuals are known, DS2) only recovery times are known, and DS3) only infection times are known. Each symbol represents the average posterior SD over 50 simulated data replicates with the error bar denoting 95% of the stochastic variation about this value, *i.e.* 95% of posterior SDs lie within the interval (note, they *do not* represent posterior credible intervals, as in Fig 4). The black line indicates a slope of −½ and the dashed black and purple dash lines indicate the sample size required for identifying a SNP with effect size 0.4 for the trait under consideration (see main text for further explanation). Parameter values are given in Eq.(10).

Instead of varying *G*_*size*_ and fixing the number of contact groups *N*_*group*_, we now fix *G*_*size*_=10 and vary *N*_*group*_. Results for this are shown in Fig 6 (represented by the crosses). This reveals a similar behaviour as seen before for the SD in *a*_*g*_ and *a*_*r*_, but crucially we find the SD in the infectivity SNP effect *a*_*f*_ now also scales with the familiar line of slope −½. The reason for this behaviour lies in the fact that infectivity is an indirect genetic effect, *i.e.* an individual’s infectivity SNP affects the disease phenotype of group members rather than its own disease phenotype [48–50]. More intuitively, this can be explained as follows. Susceptibility and recoverability SNPs of an individual directly affect its own measured disease phenotype (the former affecting its infection time and the latter affecting its recovery time). Therefore the information on which these two quantities can be inferred is expected to scale with the total number of individuals. On the other hand, as an individual’s infectivity SNP acts on all susceptible individuals sharing the same contact group, it affects the epidemic dynamics as a whole. In fact much of the information regarding infectivity comes from the overall speed of epidemics. For example, if those contact groups containing individuals with more *A* alleles consistently experience epidemics which are faster than those with fewer *A* alleles, this provides evidence that the *A* allele confers greater infectivity than the *B* allele (the situation is further complicated by the fact that differences in susceptibility can also cause this behaviour, however the algorithm can independently estimate *a*_*g*_, so removing this potential confounding). Because information about the infectivity SNP effect comes from epidemic-wide behaviour, it is expected to scale linearly with the number of contact groups *N*_*group*_ (Fig 6b), but not with the number of individuals per contact group *G*_*size*_ (Fig 5b).

**Fig 6.**
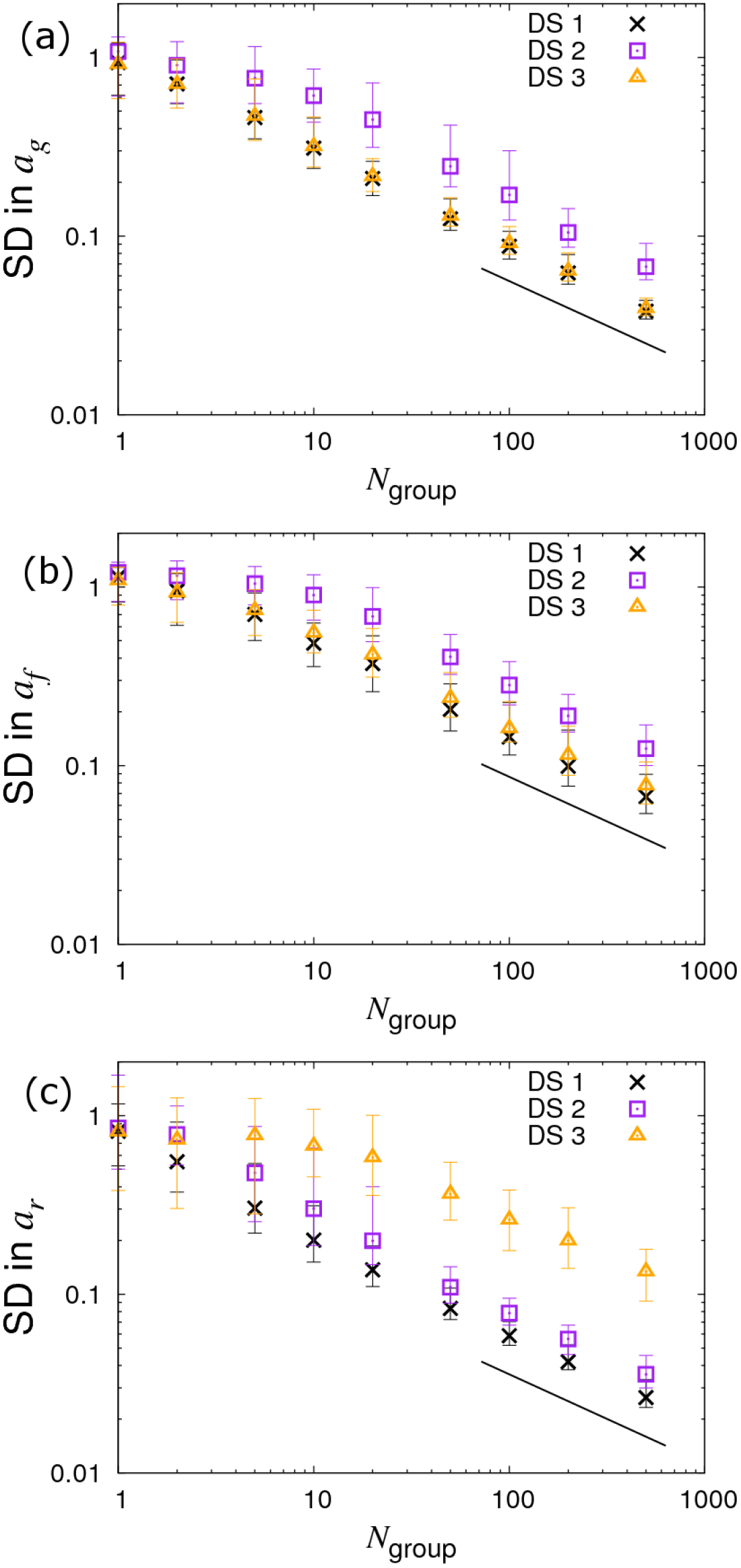
Variation in precision of the SNP effect estimates with number of groups *N*_group._ Posterior standard deviations (SDs) in SNP effects for (a) susceptibility *a*_*g*_, (b) infectivity *a*_*f*_ and (c) recoverability *a*_*r*_ from simulated data with *N*_*group*_ contact groups (which is varied) each containing *G*_*size*_=10 individuals. Different symbols represent different data scenarios: DS1) Both the infection and recovery times for individuals are known, DS2) only recovery times are known, and DS3) only infection times are known. Each symbol represents the average posterior SD over 50 simulated data replicates with the error bar denoting 95% of the stochastic variation about this value. The black line indicates a slope of −½. Parameter values are given in Eq.(10).

Finally, we investigate the case in which we fix the total number of individuals to *G*_*size*_×*N*_*group*_=1000 whilst simultaneously varying *G*_*size*_ and *N*_*group*_, as shown in Fig 7 (see crosses). In Fig 7a we find very little variation in the precision of *a*_*g*_. Interestingly, the results in Fig 7b clearly demonstrate that *larger* numbers of contact groups containing *fewer* individuals help to reduce the SD in the infectivity SNP effect *a*_*f*_. In the case of *G*_*size*_=2 the posterior SDs in *a*_*g*_ and *a*_*f*_ are actually the same due to the symmetry of this particular setup (*i.e.* each group consists of exactly one infected and one susceptible individual). Lastly, Fig 7c shows that the SD in *a*_*r*_ is largely independent of *G*_*size*_. This is because recovery is solely an individual-based process, and so happens independently of others sharing the same contact group (although in cases in which R_0_ is small, differences may result from variation in the fraction of individuals which actually become infected).

**Fig 7.**
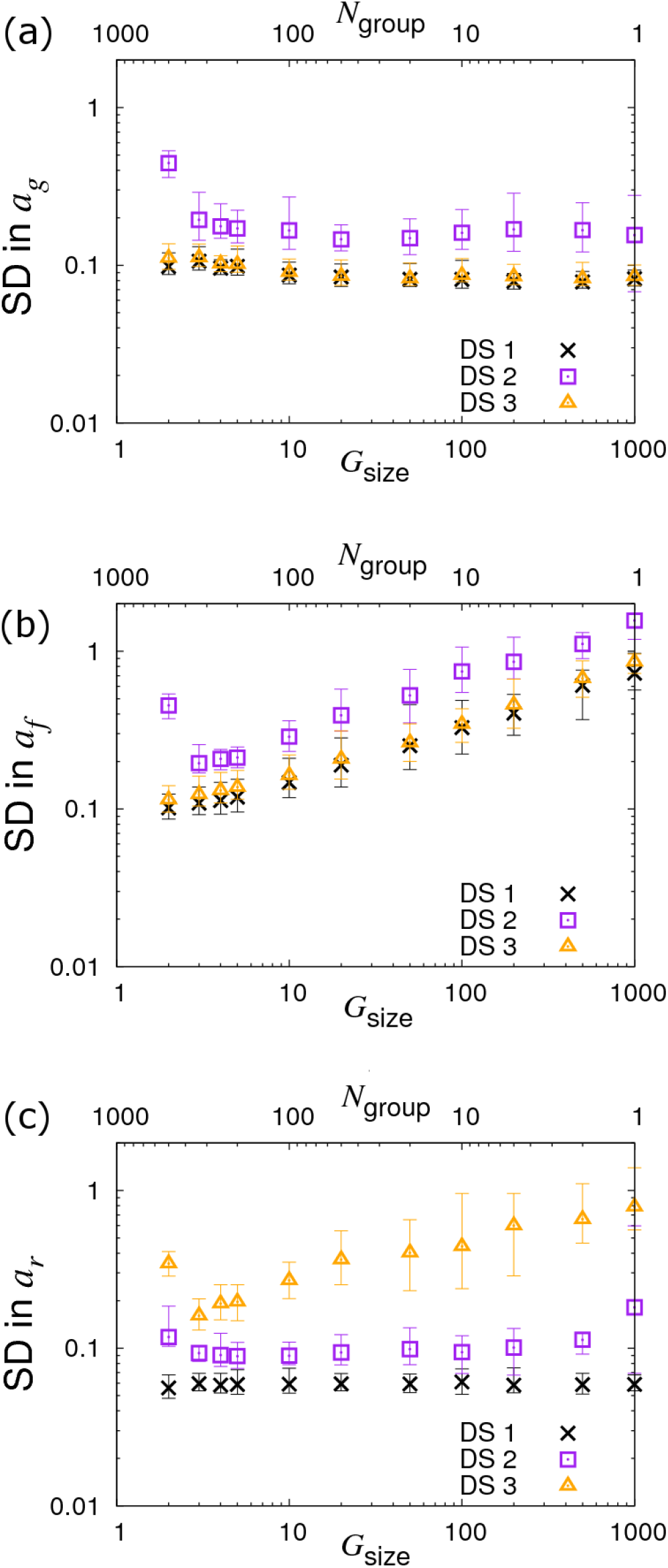
Variation in precision of the SNP effect estimates with partitioning into groups. Posterior standard deviations (SDs) in SNP effects for (a) susceptibility *a*_*g*_, (b) infectivity *a*_*f*_ and (c) recoverability *a*_*r*_ from simulated data with *N*_*group*_ contact groups each containing *G*_*size*_ individuals, both of which are varied such that the total population *N*_*group*_×*G*_*size*_ is fixed to 1000. Different symbols represent different data scenarios: DS1) Both the infection and recovery times for individuals are known, DS2) only recovery times are known, and DS3) only infection times are known. Each symbol represents the average posterior SD over 50 simulated data replicates with the error bar denoting 95% of the stochastic variation about this value. Parameter values are given in Eq.(10).

### 3.4 Dependence on allele frequency

So far we have assumed a fixed *A* allele frequency *p=*0.3 in the population. Fig 8 demonstrates what happens when this is no longer the case by varying *p*, which in turn changes the Hardy-Weinberg equilibrium frequencies for the three genotypes. We find that the curves are symmetric around a minimum of *p=*0.5 and remain remarkably flat over a large region. They only increase substantially when the minor allele frequency drops below around 10%. This result shows that the statistical power to establish SNP effects dramatically reduces when they are rare, which is consistent with observations from conventional GWAS analyses [51].

**Fig 8.**
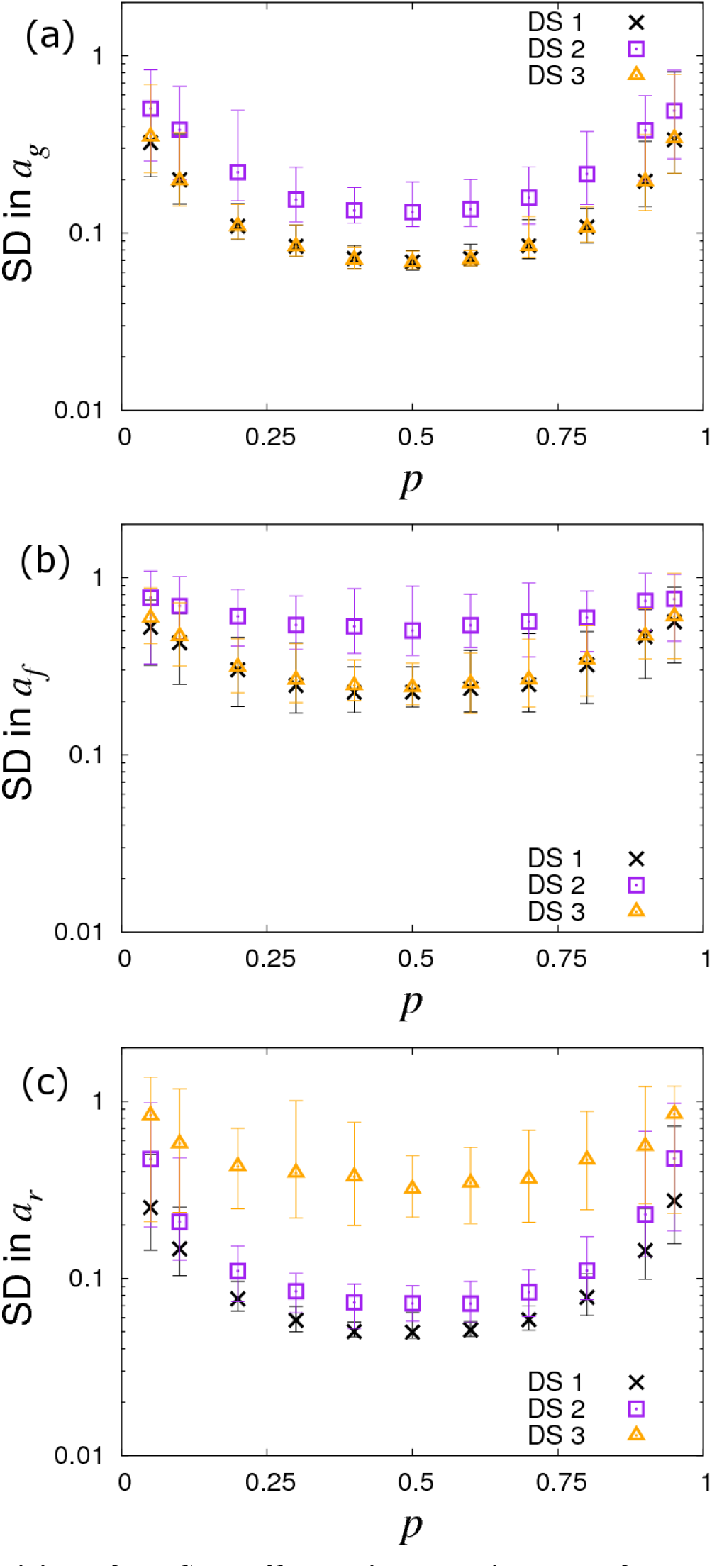
Variation in precision of the SNP effect estimates with allele frequency *p*. Posterior standard deviations (SDs) in SNP effects for (a) susceptibility *a*_*g*_, (b) infectivity *a*_*f*_ and (c) recoverability *a*_*r*_ from simulated data with *N*_*group*_=20 contact groups each containing *G*_*size*_=50 individuals. Different symbols represent different data scenarios: DS1) Both the infection and recovery times for individuals are known, DS2) only recovery times are known, and DS3) only infection times are known. Each symbol represents the average posterior SD over 50 simulated data replicates with the error bar denoting 95% of the stochastic variation about this value. Parameters used are given in Eq.(10).

### 3.5 Different data scenarios

This section shows results from the various data scenarios introduce in section 2.4, in which the infection and recovery times of all individuals are not known precisely:

#### DS2: Only recovery times known

Since *a*_*g*_ and *a*_*f*_ relate to the infection process, naïvely it might be expected that because infection times are unknown then nothing can be inferred about these SNP effects. This section, however, clearly demonstrates this not to be the case. The reason lies in the fact that whilst infection times are latent variables, the distribution from which they are sampled *is* informed by the available recovery data through the likelihood in Eq.(6).

The square symbols in Fig 5a denote the posterior SDs in the susceptibility SNP effect *a*_*g*_ under DS2. Compared to the best case scenario DS1, the SD in *a*_*g*_ increases as a result of having to infer probable infection times for individuals (as opposed to knowing them exactly). The number of individuals per group needed to identify an association for a susceptibility SNP effect of *a*_*g*_ =0.4 is now *G*_*size*_=80 (see dashed purple line in Fig 5a), as opposed to *G*_*size*_=20 in the case of DS1. Consequently to achieve an equivalent precision for *a*_*g*_ under DS2 requires around 4 times as many individuals. In the case of the infectivity SNP effect *a*_*f*_, this factor becomes approximately 4.2 (see Fig 6b, assuming a large number of contact groups), and for the recoverability it is 1.9 (see Fig 5c). These factors were found to be remarkably consistent across a broad range of group numbers and sizes (results not shown).

Estimates of prediction accuracies and bias for the case of DS2 were obtained as described in section 3.2, and results are presented in Appendix H. Compared to DS1 (Fig 4 and Table 1), The prediction accuracies tend to be slightly lower (but still above 0.5 in the majority of cases and above 0.9 for some parameters) and the bias slightly higher, reflecting the reduction in data. However, similar patterns with regards to which parameters are associated with lower prediction accuracy and bias emerge as was seen for DS1 (Fig 4).

In summary our analysis of DS2 clearly demonstrates that even when infection times are unknown, accurate inference regarding all SNP effects can be made, given sufficient data.

#### DS3: Only infection times known

The triangles in Figs 5, 6 and 7 show results under DS3 for different group sizes and group compositions. Here the SDs in the SNP effects for susceptibility *a*_*g*_ and infectivity *a*_*f*_ are found to be almost the same as for DS1 (because uncertainty in recovery times only has a very weak impact on uncertainty in the infection process). However the SD for the recovery SNP effect *a*_*r*_ is much larger, meaning that little can be inferred regarding SNP-based differences in recoverability. This is because under DS3 the only indirect information regarding recovery times comes from the very early stages of each epidemic (*e.g.* we know that the first infected individual cannot recover before the second individual becomes infected). This explains why SDs for recovery SNP effects decrease at a rate of −½ (on the log-scale) as the number of contact groups *N*_*group*_ increases (*i.e.* the triangles in Fig 6c scale with the black line) but not when the number of individuals per contact group *G*_*size*_ is changed (see Fig 5c).

#### DS4: Disease status periodically checked

Fig 9 shows results under DS4 assuming a time interval between checks of Δt. When Δt=0 (as shown on the left of this figure) the DS4 results are the same as in DS1 (because here infection and recovery times are effectively exactly known). On the other hand as checking becomes less and less frequent, the SDs in the SNP effects rise. A surprising feature is that this reduction in statistical power is perhaps less than might be expected. The vertical lines in Fig 9 represent two key timescales: 〈*t*_*I*_〉 is the average infection time as measured from the beginning of the epidemic and 〈*t*_*R*_〉 is the average recovery time (these quantities are found by averaging over a large number of simulated replicates). We see that statistical power only marginally reduces even when disease diagnostic checking is performed on a similar timescale as the epidemics as a whole. This means that results assuming DS1 which are either analytical (explored in a follow up paper [46]) or numerical, as looked at in sections 3.1–3.4, remain relevant in realistic data scenarios.

**Fig 9.**
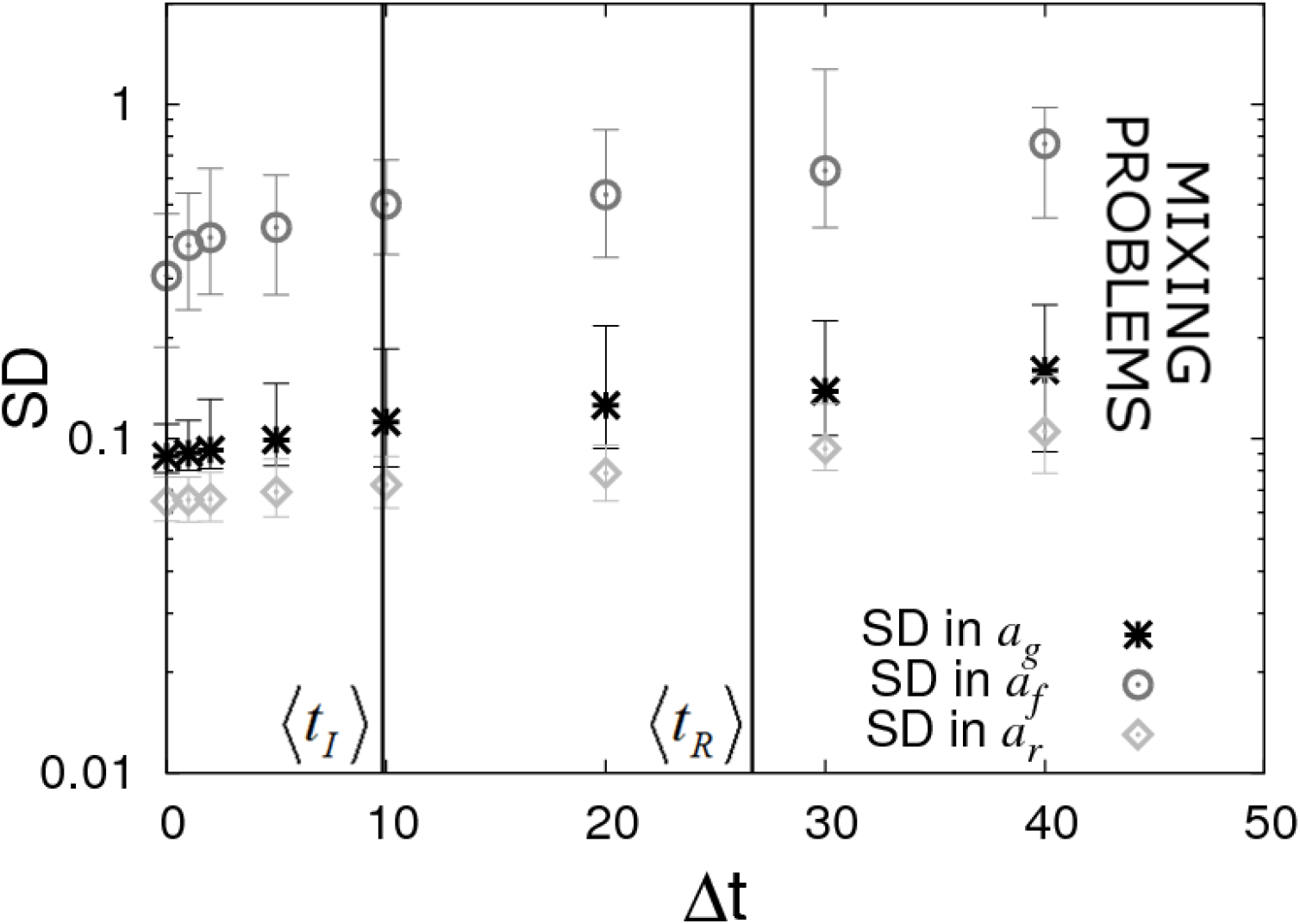
Periodic checking of disease status (DS4). Posterior standard deviations (SDs) in estimated SNP effects *a*_*g*_, *a*_*f*_ and *a*_*r*_ from simulated data with *N*_*group*_=20 contact groups each containing *G*_*size*_=50 individuals. Here it is assumed that the disease status of individuals is periodically checked with time interval Δt. Each symbol represents the average posterior SD over 50 simulated data replicates with the error bar denoting 95% of the stochastic variation about this value (with the checking times randomly shifted across these replicates) with the error bar denoting stochastic variation in posterior mean. The vertical lines represent key epidemic times: 〈*t*_*I*_〉 is the mean infection time (as averaged over an large number of simulations) and 〈*t*_*R*_〉 the mean recovery time. Parameter values given in Eq.(10).

The limit on the right hand side of Fig 9 shows the situation in which there is no information regarding infection and recovery times (*i.e.* only the initial and final states of the epidemic are observed). Unfortunately it was found to be difficult to probe this regime using SIRE due to mixing problems arising in the MCMC algorithm [52] (principally because the number of possible parameter sets and event sequences consistent with a given final outcome is vast).

The results here emphasise the fact that even relatively infrequent disease status checks provide useful data from which accurate inferences regarding SNP effects can be drawn.

#### DS5: Time censored data

In Fig 10a it is assumed the infection and recovery times are exactly known but only up to some final time *t*_*end*_ (subsequent to which no further data is available). We find that very little information is lost when restricting *t*_*end*_ to around the average recovery time 〈*t*_*R*_〉. This is largely because most individuals recover before 〈*t*_*R*_〉 as a consequence of a small number of individuals having very low recoverability (which itself arises because of the large residual variance Σ_*rr*_=1 assumed here). Given that 〈*t*_*R*_〉 is usually substantially less than the total epidemic time, from a practical point of view terminating disease transmission experiments prior to the end of the epidemic when no new infections occur, (and perhaps performing further replicates) may be beneficial. However, the effectiveness of this approach would depend on a large assumed variation in recoverability in the population, which *a priori* may be unknown.

**Fig 10.**
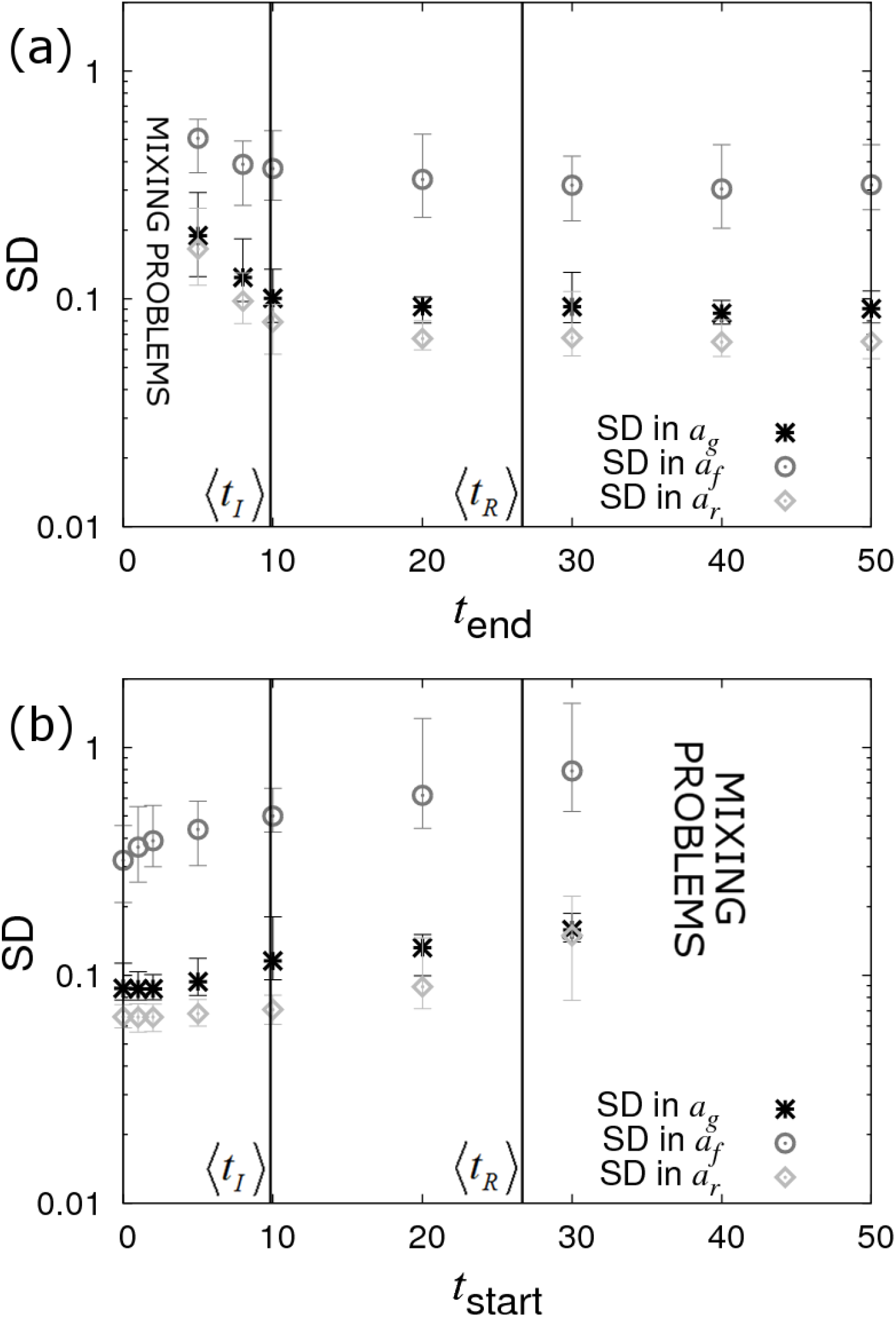
Censoring of data (DS5). Posterior standard deviations (SDs) in SNP effects *a*_*g*_, *a*_*f*_ and *a*_*r*_ from simulated data with *N*_*group*_=20 contact groups each containing *G*_*size*_=50 individuals. Each symbol represents the average posterior SD over 50 simulated data replicates with the error bar denoting 95% of the stochastic variation about this value. (a) Contact groups are observed until time *t_end_*, after which no further data is taken. (b) Contact groups are observed from time *t*_*start*_ until the end of all epidemics. The vertical lines represent key epidemic times: 〈*t*_*I*_〉 is the mean infection time (as averaged over an large number of simulations) and 〈*t*_*R*_〉 the mean recovery time. Parameter values given in Eq.(10).

Fig 10b shows the opposite scenario, in which contact groups are observed from an initial starting time *t*_*start*_ after the start of the epidemic up until its termination. This scenario may apply to field outbreaks, where sampling occurs only after notification of the outbreak. Here again we see a reduction in statistical power with increasing *t*_*start*_, but this reduction is not substantial until around the average infection time. This result is surprising, but it turns out that whilst none of the events before *t*_*start*_ are actually measured (which may include a large proportion of the total number of infection events), the disease status of all the individuals at *t*_*start*_ can be accurately inferred (because the final state is known and all the subsequent events from *t*_*start*_ are also known, the state at *t*_*start*_ is exactly specified) and this encapsulates almost the same amount of information as when the event times are precisely known.

#### General data scenario

It should be noted that the data scenarios DS1-5 considered are not comprehensive. Any combination of infection time, recovery time, disease status data and diagnostic test results can be used as inputs into SIRE. Furthermore SIRE accounts for additional uncertainties in cases in which data is missing on some individuals and where diagnostic tests are imperfect.

## 4 Discussion

The availability of dense genome-wide SNP panels has revolutionized human medicine and has paved the way for genetic disease control in agriculture. With declining genotyping costs, discovery of new disease susceptibility loci has increased exponentially over recent years, and evidence for their effective utilization in personalized medicine and livestock and plant breeding programmes continues to emerge [53–56]. However, there is increasing awareness amongst researchers and policy makers that disease susceptibility is not the only host genetic trait controlling disease incidence and prevalence in populations, and in particular that host genetic infectivity and recoverability may also constitute important improvement targets for reducing disease spread [19–22, 57, 58]. Yet, genetic loci associated with host recoverability reported in the literature are sparse, and to the best of our knowledge no infectivity SNP has yet been identified. This, perhaps, is unsurprising given that phenotypic measurements of recoverability and infectivity, such as individuals’ recovery or pathogen shedding rates are rarely available in practice and statistical inference methods to accurately infer these from available epidemic data are still in their infancy. In line with the lack of suitable statistical methods, little is known about what type and number of measurements are needed to produce unbiased and precise estimates of SNP effects for these ‘new’ epidemiological host trait phenotypes.

In this paper we developed a Bayesian methodology to allow simultaneous estimation of SNP effects for host susceptibility, recoverability and infectivity from temporal epidemic data. This methodology was validated with data from simulated epidemics, which were also used to assess how different parameter values and data scenarios representing different recording schemes in field or experimental studies may affect the estimates of SNP effects and other parameters influencing transmission dynamics. The sophisticated Bayesian algorithm outlined in this paper has been implemented into a user-friendly software tool called SIRE, which allows computationally efficient analyses to be performed by anyone with relevant epidemiological data (as shown in Appendix G, outputs typically take a few minutes of CPU time per 1000 individuals).

Our results indicate that it is possible to obtain simultaneous unbiased estimates of SNP effects for all three epidemiological host traits, in addition to that of other fixed or random effects influencing disease transmission, from temporal epidemic data. Across simulated data scenarios we found that recoverability SNP effects are generally (with few exceptions) easiest to identify, followed by susceptibility and then infectivity SNP effects. In the latter case a large number of contact groups with few individuals provide much more information than the reverse. Simulations of different data scenarios representing optimal (perfect and complete data) and practically feasible recording schemes produced the following relevant insights: firstly, even when only recovery (or death) times of individuals are known inference of SNP effects is still possible, albeit requiring around four times as many individuals to gain equivalent precision as for perfect data. Secondly, only knowing infection times marginally reduces statistical power to detect SNP effects for susceptibility and infectivity, but recovery SNP effects become difficult to detect. Thirdly, when data consists of periodic measurements of individuals’ disease status it was found that even relatively infrequent measurements (*e.g.* on a similar timescale as the entire epidemic) yields SNP effects with high precision, given sufficient data. Lastly, precise estimates of SNP effects could still be obtained with censored epidemic data.

For model validation, we chose a complex inter-dependence structure for the model parameters by assuming that the SNP under consideration is associated with all three epidemiological host traits (*i.e.* pleiotropy), but with different allele substitution effects and different modes of dominance. Furthermore, we assumed that the traits are also influenced by other fixed effects, have large residual variance (introducing much noise into the system) and are correlated, and that environmental group effects influence the within-group transmission dynamics. This choice represents an extremely challenging system in which to estimate SNP effects and in practice most real world examples are likely to be considerably less challenging as simpler structures and reduced variation/better control of variation will improve the quality of the parameter estimates.

The results from different data scenarios indicate a log-log scaling relationship with slope −½ between the precision (as measured by the SD in the posterior) of SNP effect estimates, and group size or number of groups (this relationship in analytically confirmed in a follow up paper [46]). For the majority of the simulations presented here, a moderate total population size of 1000 or less individuals was assumed. The corresponding posterior standard deviations for estimated SNP effects were generally above 0.01, and in the case of infectivity effects, more often above 0.1. This would suggest that for datasets comprising of 1000 individuals or less, SIRE is only able to detect SNPs of large effects on the epidemiological host traits, but identification of SNPs of small to moderate effects on this trait requires significantly more data, in particular for infectivity.

We chose a dataset comprising of 1000 individuals partly because of computational efficiency but also because generating datasets of this size seems feasible for transmission experiments in plants and most domestic livestock species, in particular aquaculture species [20, 59, 60]. However, many existing field data, in particular in dairy cattle populations with routine genotyping and frequent recordings of disease phenotypes *e.g.* for mastitis, bovine Tuberculosis, and other infectious diseases [61–63] already exceed this number by several orders of magnitude. As genotyping costs continue to fall and automated recording systems are applied at rapidly increasing frequency in agriculture [64, 65], the possibility of identifying SNPs with small to moderate effects on the epidemiological host traits, and their mode of dominance, which was poorly estimated for the given sample size, would appear to be well within reach in the near future.

It is widely recognised that disease traits are for the most part polygenic, *i.e.* regulated by many genes each with small effect, and hence that SNPs with large effect on disease phenotypes are the exception rather than the norm [10, 62]. This is partly due to the fact that observed disease phenotypes, such as individuals’ binary infection status or infection time are the result of many interacting biological processes, each controlled by a different set of genes or genetic pathways and characteristics of the wider population. Hence the impact of an individual gene on the disease phenotype is diluted. In contrast, the relative impact of a particular gene on traits that are more closely related to specific biological processes, such as *e.g.* pathogen entry, replication or shedding affecting susceptibility, recoverability or infectivity, respectively, may be higher [66]. Therefore, it is not unreasonable to assume that SNPs with moderate to large effects on these epidemiological traits, and in particular on host infectivity, may indeed exist. Evolutionary theory suggests that alleles that confer low susceptibility to infection or fast recoverability from infection are subject to strong directional selection when individuals are commonly exposed to infection [67]. Hence, such beneficial alleles tend to become fixed within only a few generations, and consequently, SNPs with large effects on disease susceptibility or recoverability would be expected to occur primarily only in populations that have not experienced strong selection pressure for these traits. This is exemplified in the case of Infectious Pancreatic Necrosis (IPN) in farmed Atlantic salmon that have only undergone a few generations of selection, where a single SNP explains most of the variation in mortality of fish exposed to the IPN virus [60, 68]. In contrast, selection pressure on infectivity is expected to be relatively low, since an individual’s infectivity genes affect the disease phenotype of group members rather than its own disease phenotype [34, 48, 69]. Therefore, infectivity SNPs with large effect may indeed exist, and may now be identifiable with the methods presented here.

The approach developed in this study and integrated into SIRE complement and succeed previous studies that aimed to develop statistical methods for estimating genetic effects for the different host epidemiological traits [25, 30–32, 34]. The key novelty of our approach lies in its ability to estimate genetic and non-genetic effects associated with all three epidemiological host traits from a range of temporal epidemic data, even when that data is incomplete.

### Applications

Many disease challenge experiments and field studies have identified SNPs with moderate to large effects on measurable disease resistance phenotypes [54, 55, 70]. However, the role of these SNPs on transmission dynamics is often poorly understood. For example, it is generally not known whether individuals that carry the beneficial allele for *e.g.* surviving infectious challenge are less likely to become infected (*i.e.* less susceptible), or more prone to surviving infection (*e.g.* due to better recoverability), and also less prone to transmitting infection, once infected (*i.e.* less infective). From an epidemiological perspective, SNPs with favourable pleiotropic effects on all three host epidemiological traits are highly desirable for preventing or mitigating disease spread [71]. In contrast alleles associated with better survival in existing GWAS would only bring the expected epidemiological benefits if they don’t simultaneously confer greater infectivity. In other words, knowing the SNP effects for all three underlying epidemiological host traits is desirable for effective employment of genetic disease control. Based on the results in this paper, SIRE can readily be applied to disentangle such SNP effects using data from transmission experiments or field studies. Furthermore, although this paper focused on estimating SNP effects, SIRE could also immediately be applied to estimating breed, age, sex, treatment or vaccination effects, or any other factor that may affect disease spread, even if genetic information is absent.

### Limitations of the current approach and future work

One of the potential practical limitations for accurately estimating infectivity SNP effects is that they require a large number of epidemic groups. Previous work has shown that experimental designs can have a significant impact on the precision and accuracy with which model parameters can be estimated (as demonstrated to some extent in this paper and also investigated for indirect genetic effects in numerous other studies [49]). In particular, theoretical studies indicate that significant improvement in estimates of infectivity effects can be achieved by appropriately grouping genetically related individuals [32, 34]. Whilst this paper focused entirely on a fixed *A* allele frequency *p* across groups, a follow up paper [46] will show that appropriate variation in genotypes within and across contact groups can lead to substantial improvements in the precision of the infectivity SNP effect *a*_*f*_, without the need for large numbers of epidemic contact groups (interestingly, the susceptibility and recoverability SNP effects cannot be substantially improved in this way).

A tool such as SIRE that can accurately estimate the effects of single SNPs on hitherto inaccessible epidemiological traits presents an important first step towards creating a statistically consistent scheme for performing GWAS on epidemiological traits using potentially incomplete data. GWAS, however, typically contains additional features beyond the scope of the simple single SNP analysis presented here. In particular, the current software focuses on one SNP at a time for estimating genetic effects for susceptibility, infectivity and recovery, but ignores the contributions of other genes on these traits. In the current model design these are incorporated into the residual effects. This simplifying assumption may have little impact for appropriately designed transmission experiments, but may lead to biased estimates of SNP effects if genetically similar individuals are not randomly distributed across groups. Theory also suggests that the required sample size for GWAS increases with the number of loci affecting the trait under consideration [72]. Hence, further model development is required for enabling GWAS for the three underlying epidemiological host traits. Previous work in our group developed a Bayesian algorithm for estimating polygenic effects for host susceptibility and infectivity from incomplete epidemic data [31]. Combining both approaches may prove a useful way forward to allow estimation of genetic effects under all realistic genetic architectures and population structures.

In summary, this paper introduces, for the first time, software that can estimate genetic and non-genetic effects for susceptibility, infectivity and recoverability simultaneously. This user-friendly tool can be applied to a range of experimental and field data and will help move genetic disease control significantly forward, beyond the focus on genetic improvement of resistance alone.

## APPENDIX A: Recovery dynamics

Fig A shows examples of gamma distributions for three different values of shape parameter *k*=1, 2 and *10*. A common approach when modelling epidemics is to assume *k*=1, corresponding to the Markovian assumption of constant recovery probability. From a biological point of view, however, this is rather unrealistic. Typically after infection, the host’s immune system takes some time to respond, *e.g.* to generate the appropriate antibodies to fight off the infection. Thus, for many real diseases *k* may be large.

**Fig A.**
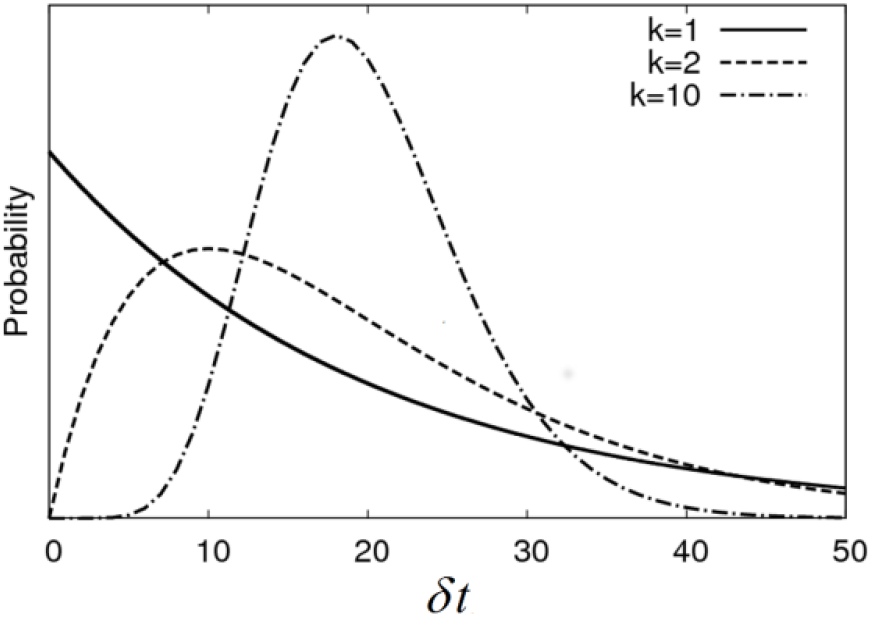
Infection duration. Shows the probability distribution for an individual to recover from their infection a time *δt* after they were infected.

Consequently, incorporation of a gamma distributed infection duration (which is characterised by two parameters, a mean and a shape parameter, instead of just one for the exponential distribution) allows the model to more realistically capture true disease dynamics.

## APPENDIX B: Derivation of the likelihood

The likelihood in Eq.(6) represents the probability the model in section 2.1 generates a certain set of events ξ, assuming a given set of parameters θ. This can be calculated by multiplying the probabilities for each of the individual sampling steps used in the simulation procedure, as described in Appendix F.

During initialisation of a given contact group, the infection duration for the initially infected individual *j* is sampled with probability

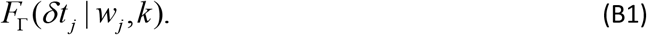

We now consider each event *e* in turn. In step 1 of Appendix F, the inter-event time Δt is sampled from the exponential distribution 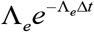 (where Λ_*e*_(*t*_*e*_) is the total infection rate Eq.(7) evaluated immediately prior to *t*_*e*_, the time of event *e*). Two possibilities exist for event *e*:

1. **It is a recovery event**. Considering the algorithm in Appendix F, step 2(a) would have been branched to, and this only happens if the sampled value for Δt is greater than the observed inter-event time *t*_*e*_−*t*_*e−1*_. The probability of this is calculated from the following integral:

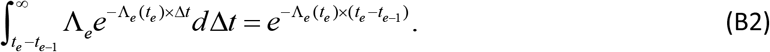
2. **It is an infection event**. Again, considering the algorithm in Appendix F, step 2(b) would have been branched to, and this happens with probability 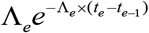 (which comes from the exponential distribution above). The individual *j* which becomes infected is selected with probability *λ*_*j*_/Λ_*e*_. Finally, the infection duration of that individual *δt*_*j*_ is sampled from a gamma distribution with probability density function *F*_Γ_(*δt*_*j*_ | *w*_*j*_, *k*). Combining these three contributions gives an overall probability:

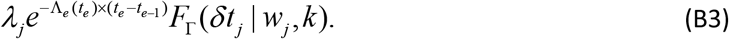

Multiplying the results from Eqs.(B1), (B2) and (B3) for all the infection and recovery events leads to the likelihood for a single contact group of:

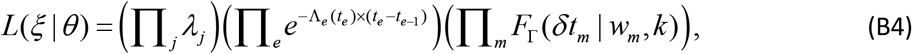

where *j* goes over individuals that become infected (*excluding* those which initiate epidemics), *m* also goes over individuals that become infected but *including* those which initiate epidemics and *e* goes over both infection and recovery events (with corresponding event times *t*_*e*_).

Since contact groups are assumed to be independent, the likelihood for multiple contact groups is simply the product of each separate one, as shown in Eq.(6).

## APPENDIX C: Prior definitions for model parameters

The default prior in SIRE (which can be altered), which is used for all the results in this paper, is largely uninformative but does place upper and lower bounds on many of the key parameters to stop them straying into biologically unrealistic regimes. Bounding parameters in this way was found to be especially important when considering relatively uninformative data scenarios when unbounded flat priors could lead to improper posterior probability distributions. Specifically, a uniform prior between −2.3 and 2.3 was chosen for *a*_*g*_. This corresponds to assuming that it is biologically unrealistic for a single SNP to change the susceptibility of individuals by more than a factor of 100 between *AA* and *BB* individuals^1^. An identical uniform prior was also placed on *a*_*f*_, *a*_*r*_ and on each of the fixed effects in ***b*_*g*_**, ***b***_***f***_ and ***b*_*r*_**. Similarly, a uniform prior between −3.45 and 3.45 was placed on each of the residuals **ε**_***g***_, **ε**_***f***_ and **ε**_***r***_, This larger range reflects a potential factor of 1000 variation across individuals (chosen to be larger as residual contributions account for all other SNPs as well as non-genetic factors, as opposed to just the effect of the single SNP under analysis).

The scaled dominance factors Δ_*g*_, Δ_*f*_, Δ_*r*_, were chosen to have uniform priors between 1 and −1, *i.e.* going from complete dominance of *A* to complete dominance of *B* ^*2*^. The prior for the shape parameter *k* was chosen to be uniform between 1 and 10, where 1 represents a Poisson random process and 10 represents a situation in which recovery times of individuals are highly concentrated around their mean.

Because parameters β and γ depend on the timescale over which the epidemic is measured^3^, which is partly pathogen specific, placing informative priors on β and γ by default is not appropriate (although it can be done). Instead a uniform prior between 0 and 20 was placed on the equivalent basic reproductive ratio R_0_ ^4^, which *is* a dimensionless quantity.

## Appendix D: MCMC procedure

Markov chain Monte Carlo (MCMC) produces a list of (correlated) parameter θ^*q*^ and event time ξ^*q*^ samples drawn from the posterior probability distribution in Eq.(5), where *q* indexes sample number.

1. **Initialisation** – MCMC is initially started with some set of parameters θ^1^ and event times ξ^1^ consistent with the data *y*. In the case of parameters, θ^1^ is simply sampled from the prior distribution π(θ). For events, initialisation depends on the available data: DS1) events are defined by the data, DS2) infection times are exponential sampled backwards in time from the first observed recovery time^5^, DS3) recovery times are exponentially sampled forwards in time from the final observed infection time^6^, DS4) infection and recovery times are sampled uniformly in the time intervals identified between successive diagnostic tests, and DS5) events within the observation period are defined by the data, infection events prior to this time period are exponentially sampled backward in time and similarly recoveries events after this time period are exponentially sampled forward in time.
2. **Iteration** - MCMC operates by proposing changes to the model parameters θ and event times ξ and accepting or rejecting these changes in accordance with a Metropolis-Hastings probability. In this way the underlying dynamics are able to explore all potential possibilities consistent with the observations.

A single MCMC “update” consists of making the following sets of proposals:

#### MCMC UPDATE

**Parameters –** Each individual parameter in θ=(*β*, *γ*, *k*, *a*_*g*_, *a*_*f*_, *a*_*r*_, Δ_*g*_, Δ_*f*_, Δ_*r*_, ***b***_***g***_, ***b***_***f***_, ***b***_***r***_, **ε**_***g***_, **ε**_***f***_, **ε**_***r***_,, **Σ**, ***G***, σ_G_, denoted by θ_*j*_, is considered in turn. A proposed value is drawn from a normal distribution centred on the parameter’s current chain value

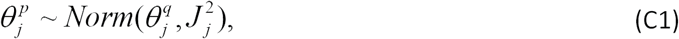

with all other parameters in θ^*p*^ remaining the same as in θ^*q*^ (if this produces an inconsistent value, e.g. *β* becomes negative, the proposal is immediately rejected). The proposal is accepted with Metropolis-Hastings probability

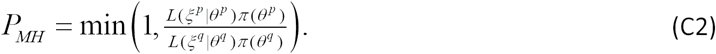

If accepted, we set θ^*q+1*^= θ^*p*^ else θ^*q+1*^= θ^*q*^ with ξ^*q+1*^= ξ^*q*^.

Tuning *J*_*j*_ in Eq.(C1) is important. If it is too large, very few proposals will be accepted and if too small, mixing will be slow. Motivated by adaptive MCMC [73, 74], a robust heuristic method for optimising *J*_*j*_ within the burn-in period is as follows. Initially, *J*_*j*_ is set to a small quantity. Each time a proposed change on parameter *j* is accepted *J*_*j*_ is updated according to

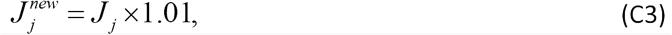

and when rejected

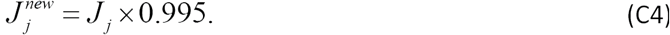

These numerical factors are chosen for two reasons: Firstly, the updates in Eqs.(C3) and (C4) balance each other out when acceptance occurs around 33% of the time, leading to a steady state solution for *J*_*j*_. Secondly, they are chosen to be sufficiently close to *1* to prevent large fluctuations in *J*_*j*_, but sufficiently far to allow the steady state solution to be found within the burn-in period.

In addition to the single parameter proposals outlined above, joint proposals are also made on residuals along with their corresponding covariance matrix, as described in Appendix E. The reason these joint updates are necessary is that **Σ** and **ε** are often highly correlated within the model (as determined by Eq.(9)). In many cases the data provides little information regarding **Σ** itself, so these correlations can lead to extremely slow mixing.

**Events –** In situations in which ξ is not precisely known (*i.e.* data scenarios other than DS1), the unknown latent event times must be stochastically changed in accordance with the posterior distribution. Each individual *j* is considered in turn. In the case of proposing changes to the infection time of *j*, the following normal distribution is sampled from:

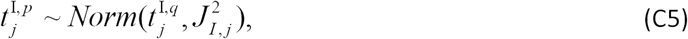

with all other event times in ξ^*p*^=(*t*^I^,*t*^R^) remaining the same as in ξ^*q*^. If this proposed infection time exceeds the recovery time 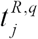 the proposal is immediately rejected (infections cannot occur after recoveries). The proposal in Eq.(C5) is accepted with probability

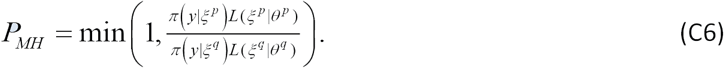

If accepted, we set ξ^*q*+1^= ξ^*p*^ else ξ^*q*+1^= ξ^*q*^ with θ^*q*+1^= θ^*q*^.

A similar proposal to Eq.(C5) is used to change recovery times. The jumping parameters *J*_*I,j*_ and *J*_*R,j*_ are again tuned to give an average acceptance probability of around 33% (using the same method as in Eqs.(C3) and (C4) above). Furthermore, there are data scenarios in which it is not possible to tell whether an individual has become infected or not. In these cases it is necessary to include additional proposals which insert infection/recovery event pairs and, conversely, other proposals which remove them.

Each of the proposals above have been optimised to calculate only those parts of the likelihood that change, *e.g.* if the infection time of individual *j* is altered then only the change in likelihood in the period between then initial and proposed times within *j*’s contact group needs to be calculated. Furthermore, when making some sets of proposals computational speed is substantially increased by pre-calculating terms in the likelihood which remain unchanged.

## Appendix E: Posterior-based proposals for residuals

This appendix describes special joint proposals in the residuals (**Σ**,**ε**) which are used to aid MCMC mixing. Specifically three type of proposal are considered: 1) those which randomly make changes to Σ_*gg*_, Σ_*g*f_, or Σ_*gr*_ and stochastically alter **ε**_**g**_ (with everything else kept fixed), 2) those which randomly make changes to Σ_*ff*_, Σ_*gf*_, or Σ_*rr*_ and stochastically alter **ε**_***f***_ (with everything else kept fixed), and 3) those which randomly makes changes to Σ_*rr*_, Σ_*gr*_, or Σ_*fr*_ and stochastically alter **ε**_***r***_ (with everything else kept fixed). Here we describe just one of these possibilities, but others can be found by suitably permuting indices.

So-called “posterior-based proposals” (PBPs) [1] consist of three steps:

**Step 1** – A proposed is made to one of the parameters in the model. In this example

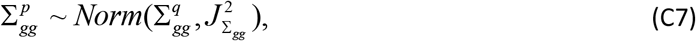

with the value of 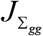 tuned to give an average acceptance probability of around 33% (using the same method as in Eqs.(C3) and (C4) above).
**Step 2** – Each element ε_*g,j*_ in the **εg** is considered in turn. The values of means *μ*_*q,j*_ and *μ*_*p,j*_ and standard deviations σ_*q,j*_ and σ_*p,j*_ are calculated such that the normal distributions they characterise approximate, to some level of accuracy, the true posterior distributions for ε_*g,j*_ in the initial and proposed states (details on how this is done are discussed below). If σ_*p,j*_>σ_*q,j*_ then for each individual *j* the proposed residuals are sampled using

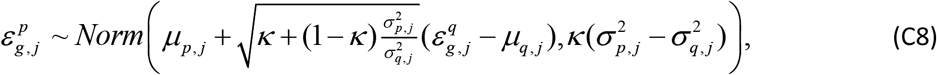

where κ is a tuneable constant set to 0.03, and when σ_*p*_≤σ_*q*_

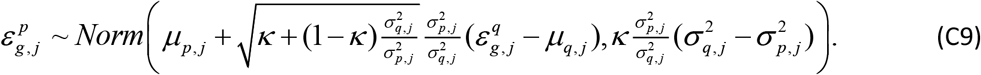 The somewhat complicated expressions in Eqs.(C8) and (C9) are specifically designed such that if the posterior *is* truly represented by the normal approximations (as characterised by *μ*_*q,j*_, *μ*_*p,j*_, σ_*q,j*_ and σ_*p,j*_), the proposal is accepted with probability 1.
**Step 3** – The proposed combination 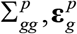 is accepted or rejected with Metropolis-Hastings probability

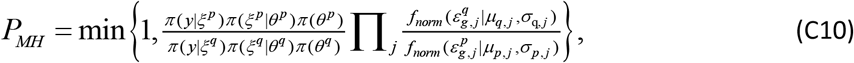

where *j* goes over all individuals and

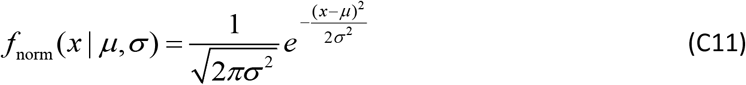

is the Gaussian probability density function.

### Optimisation

For efficient implementation of this approach it is necessary to generate *μ*_*q,j*_, *μ*_*p,j*_, σ_*q,j*_ and σ_*p,j*_ in a suitable manner. One approximation is to set them to values implied by prior in Eq.(9):

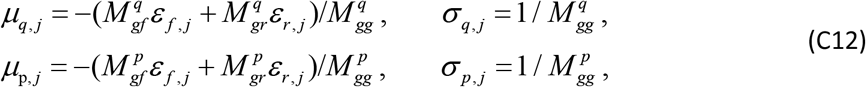

where **M** is the inverse of the covariance matrix **Σ**^**−1**^. This approximation leads to so-called “model-based proposals” (MBP). A more accurate approximation (because it takes into account the observed events, which themselves are informed by the data) is given by

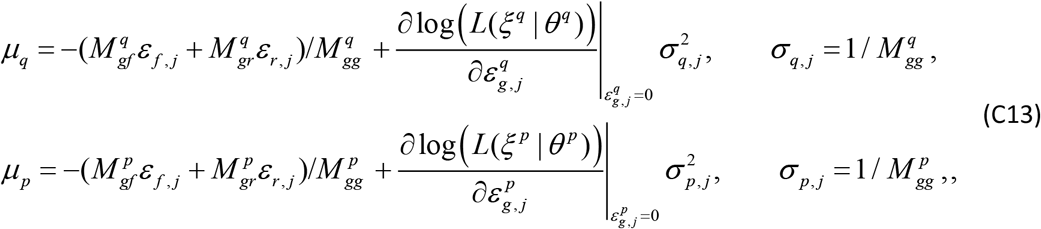

and this gives a first order “posterior-based proposal” (PBP). Although Eq.(C13) is computationally slower to calculate than Eq.(C12) (because it contains gradients in the log-likelihood), it can lead to larger jumps in parameter space compared to a corresponding MBP resulting in improved mixing. Empirically it was found that applying PBPs to parameters relating to susceptibility and recoverability and MBPs to parameters relating to infectivity was the fastest approach to take.

## Appendix F: Simulation

The Doob-Gillespie algorithm [43] provides a means of taking into account inherent stochasticity in Markovian compartmental models (*i.e.* models for which the transition rates depend solely on the current state of the system). The model used in this paper combines Markovian infection transitions with more realistic non-Markovian recovery dynamics. Below we describe how these recovery events are incorporated into the standard Doob-Gillespie framework.

The purpose of this procedure is to build up a time ordered sequence of infection and recovery event times indexed by event number *e*. The following notation is used: *t*_e_ is the event time, *x*_e_ is the event type (infection *in.* or recovery *re.*), *j*_*e*_ is the affected individual, and 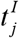 and 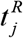 are the infection and recovery times for individual *j*, respectively.

**Initialization:** Each epidemic is assumed to be started by one (or potentially more) initially infected individual *j* at some initial time point *t*_*init*_. The infection duration *δt*_*j*_ for this individual is drawn from a gamma distribution parameterised in terms of a mean and shape parameter:

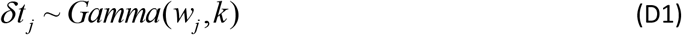

(note, the dependency of *w*_*j*_ on θ is given through Eqs. (2) and (3)). This allows us to set 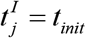 and 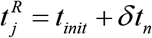. Individual *j* is then placed onto a list 𝓡, which represents all currently infected individuals. We set event index to *e*=*1*.

**Step 1**: Calculate the time to the next infection event. This is done by first evaluating the total transition rate that any individual becomes infected

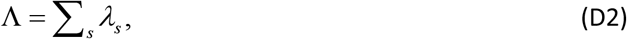

where the sum *s* goes over all currently susceptible individuals and the force of infection *λ*_*s*_ (which gives the probability per unit time of *s* becoming infected) is given by Eq.(1). In accordance with a Poisson process, the time to the next infection event is generated by drawing a sample from the exponential distribution Λ*e*^−ΛΔ*t*^. In practice, this is achieved by selecting an inter-event time using

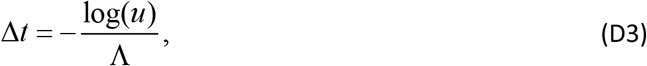

where *u* is a (uniform) randomly generated number between *0* and *1*. The new event time is then defined by

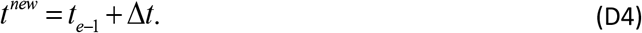

**Step 2**: Choosing the event type. For the SIR model two possibilities exist:

a. If *t*^new^ is greater than the smallest recovery time of all the individuals in 𝓡, which we label *j*_*min*_, then we remove *j*_min_ from 𝓡 and set

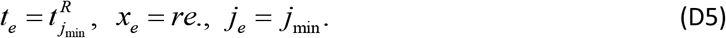
b. Otherwise, we set

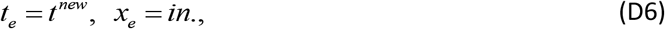

and select the actual individual that becomes infected with probability

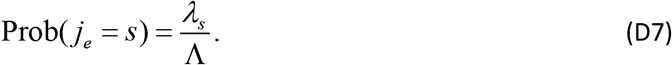

The infection duration 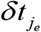 for *j*_*e*_ is sampled using Eq.(D1), and the infection and recovery times are set to

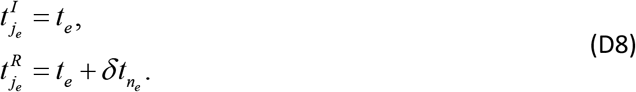

Individual *j*_*e*_ is then placed onto the list 𝓡.

**Step 3:** Increment *e* and jump to step 1 if there are any remaining infected individuals.

**End:** Insert recovery times for any remaining individuals *j* in 𝓡

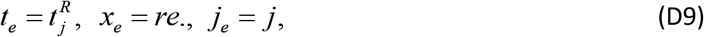

incrementing *e* after each addition.

## Appendix G: Computational speed estimate for SIRE

**Fig G.**
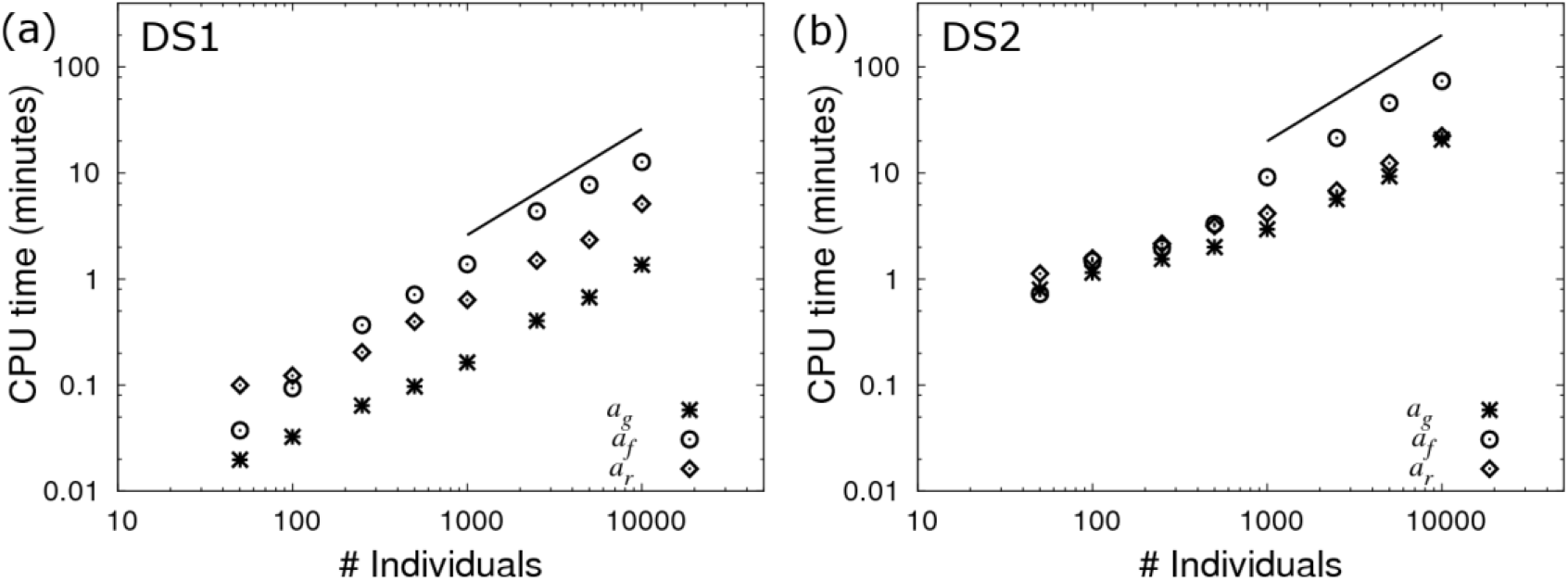
SIRE speed. The CPU time taken for SIRE to generate 100 independent posterior samples for the SNP effects as a function of the number of individuals (where the group size is taken to *G*_*size*_=50), as estimated using the effective sample size [1]. Simulated data was generated using the base parameter set in Eq.(10) with (a) known infection and recovery times (DS1) and (b) known recovery times (DS2).

Fig G shows the CPU time SIRE takes to estimate the SNP effects as a function of the total number of individuals (this is based on a single 2GHz core). We find that the SNP effect associated with infectivity takes the longest to accurately estimate. The approximate linear scaling (represented by the solid black lines) means that SIRE is expected to take around one minute per 1000 individuals to generate 100 representative samples from the posterior under DS1 and around 10 minutes for DS2.

## Appendix H: Parameter prediction accuracy under DS2

Fig H shows results when repeating the analysis is section 3.2 of the paper, but this time taking the scenario in which infection times are unknown (*i.e.* DS2). The corresponding regression analysis on these points is presented in Table H. The average SDs in Table H are smaller than those in Table 1, reflecting the reduction in statistical power under this data scenario. A factor of two difference in SD corresponds to four times as many individuals required for equivalent accuracy.

**Fig H.**
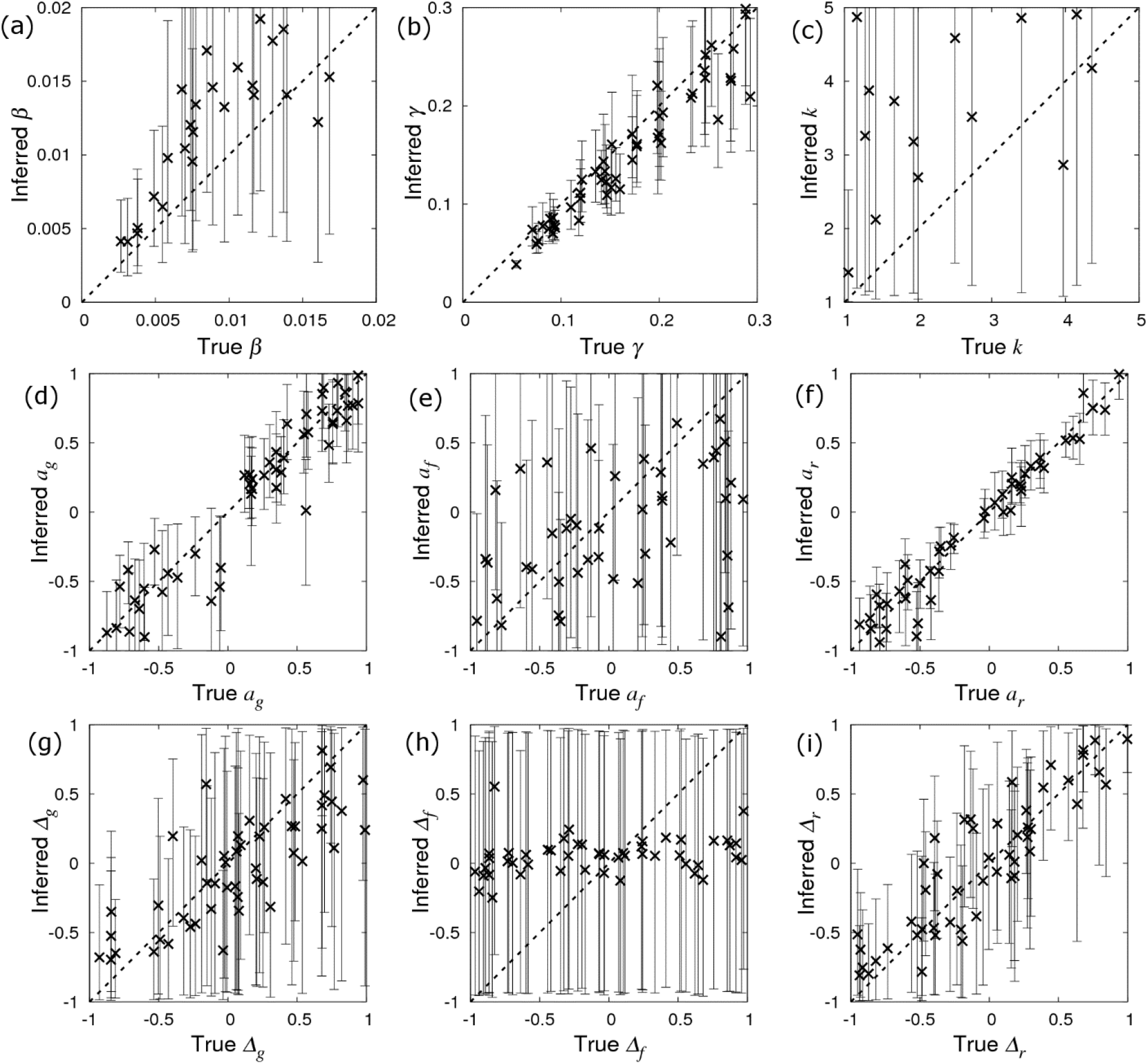
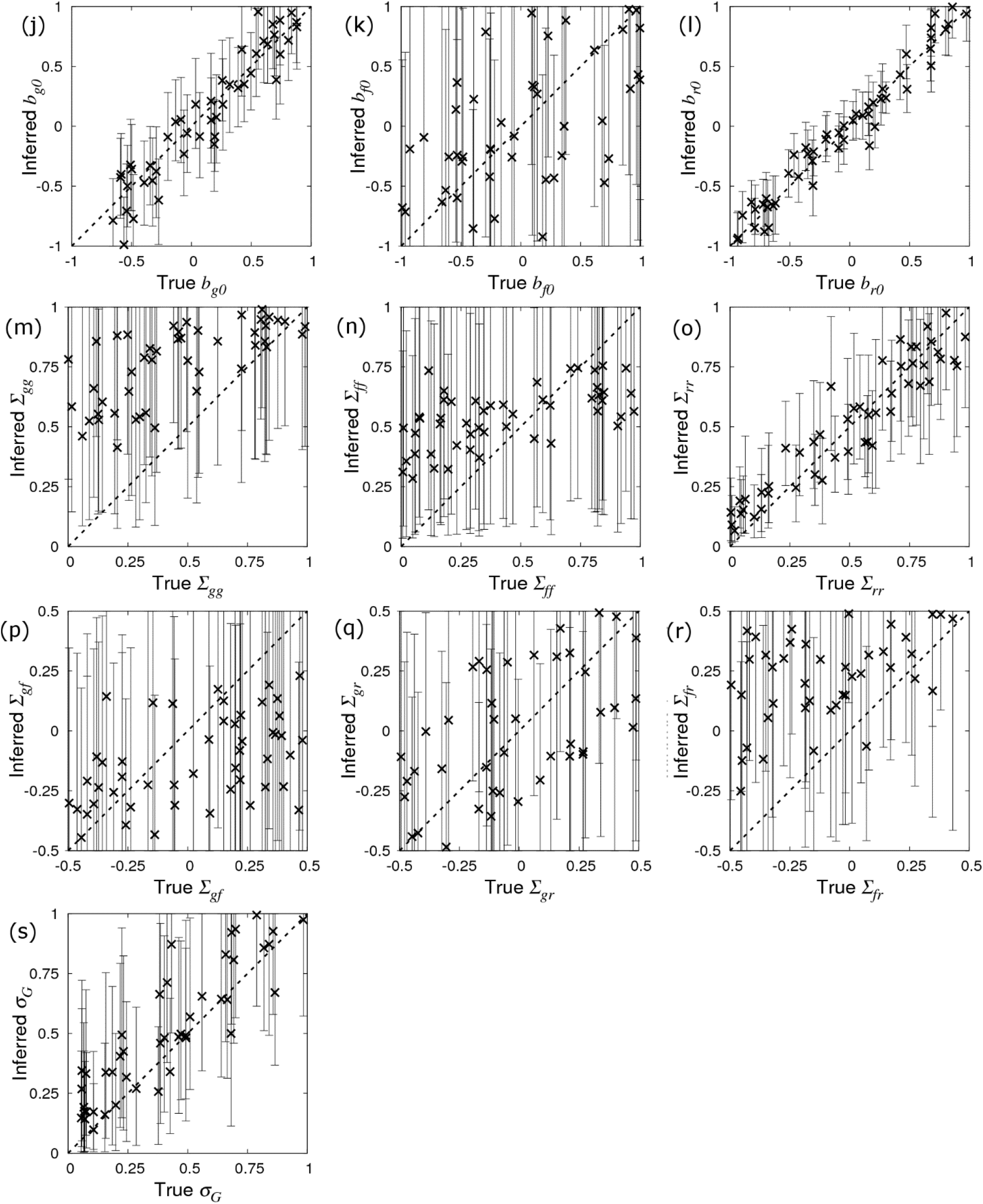
Prediction accuracy and bias. These plots summarise inferred posterior distributions for parameters compared to their true value. Simulated data was generated using the base parameter set in Eq.(10) except for a single parameter which was singled out in each of the sub-plots above*. Crosses correspond to the inferred posterior mean (with error bars indicating 95% credible intervals) of the selected parameter (whose true value is on the x-axis) when SIRE is applied to a single simulated data set consisting of recovery times (*i.e.* DS2) from *N*_*group*_=20 contact groups each containing *G*_*size*_=50 individuals. Prediction accuracies, and the intercept and slope of regression lines fitted to the data points are given in Table 1. (*Additionally for (g) *a*_*g*_=0.4, (h) *a*_*f*_=0.4 and (i) *a*_*r*_=0.4, such that dominance has an effect).

**Table H.** Prediction accuracy and bias under DS2. Prediction accuracy is defined as the correlation between the inferred and true parameter values (a value of one implies perfect inference). The *y*-intercept and slope are taken from regression lines fitted through the data point (a *y*-intercept of zero and slope of one indicates no bias). Av. SD gives the average posterior standard deviation across all datasets.

| Parameter | Accuracy | y-intercept | Slope | Av. SD | Description |
| --- | --- | --- | --- | --- | --- |
| $\beta$ | 0.831 | 0.003 | 1.191 | 0.005 | Average transmission rate |
| $\gamma$ | 0.956 | -0.008 | 0.946 | 0.024 | Average recovery rate |
| $k$ | 0.494 | 3.454 | 0.516 | 2.272 | Recovery shape parameter |
| $a_g$ | 0.950 | -0.004 | 0.995 | 0.173 | SNP effect for susceptibility |
| $a_f$ | 0.355 | -0.019 | 0.370 | 0.525 | SNP effect for infectivity |
| $a_r$ | 0.979 | -0.006 | 1.022 | 0.111 | SNP effect for recoverability |
| $\Delta_g$ | 0.791 | -0.079 | 0.619 | 0.387 | Dominance factor (per trait) |
| $\Delta_f$ | 0.260 | 0.056 | 0.054 | 0.544 | |
| $\Delta_r$ | 0.891 | -0.040 | 0.857 | 0.249 | |
| $b_{g0}$ | 0.961 | -0.033 | 1.083 | 0.188 | Fixed effect (per trait) |
| $b_{f0}$ | 0.411 | -0.004 | 0.469 | 0.647 | |
| $b_{r0}$ | 0.978 | 0.011 | 0.980 | 0.111 | |
| $\Sigma_{gg}$ | 0.723 | 0.581 | 0.416 | 0.236 | Residual covariance matrix |
| $\Sigma_{ff}$ | 0.503 | -0.141 | 0.312 | 0.330 | |
| $\Sigma_{rr}$ | 0.574 | 0.014 | 0.655 | 0.296 | |
| $\Sigma_{gf}$ | 0.588 | 0.440 | 0.233 | 0.266 | |
| $\Sigma_{gr}$ | 0.537 | 0.292 | 0.382 | 0.248 | |
| $\Sigma_{fr}$ | 0.941 | 0.101 | 0.817 | 0.113 | |
| $\sigma_G$ | 0.906 | 0.131 | 0.936 | 0.182 | SD of group effects |

## Footnotes

1 This factor comes from the exponential dependency in Eq.(1) coupled with the result e^2×2.3^ =100.

2 If over dominance is considered a possibility, this prior distribution would be extended.

3 E*.g.* if measurements are made in hours then β and γ would be very different to if they were made in days.

4 Defined by R_0_=β(〈N〉−1)/γ, where 〈N〉 is the average population size of contact groups as taken from the data, this represents the number of infections one typically infectious individual generates on average over the course of its infectious period in an otherwise uninfected population.

5 The standard deviation in recovery times is used to set the decay rate.

6 The standard deviation in infection times is used to set the decay rate.

